# Bcl11b orchestrates subcerebral projection neuron axon development via cell-autonomous, non-cell-autonomous, and subcellular mechanisms

**DOI:** 10.1101/2024.10.20.619265

**Authors:** Yasuhiro Itoh, Mollie B. Woodworth, Luciano C. Greig, Anne K. Engmann, Dustin E. Tillman, John J. Hatch, Jeffrey D. Macklis

**Affiliations:** Department of Stem Cell and Regenerative Biology, and Center for Brain Science, Harvard University, Cambridge, Massachusetts 02138, USA; Voyager Therapeutics, Lexington, MA 02421, USA; Neuroscience Program, Bates College, Lewiston, ME 04240, USA; Department of Ophthalmology, University of California, San Francisco, CA 94143, USA

## Abstract

Both cell-intrinsic competency and extracellular cues regulate axon projection, but mechanisms that coordinate these elements remain poorly understood. Subcerebral projection neurons (SCPN) extend their primary axons from cortex through subcortical structures, including the striatum, targeting the brainstem and spinal cord. We identify that the transcription factor Bcl11b/Ctip2 functions in multiple independent neuron populations to control SCPN axon development. *Bcl11b* expressed by SCPN is required cell-autonomously for axonal outgrowth and efficient entry into the internal capsule within the striatum, while *Bcl11b* expressed by medium spiny neurons (MSN) non-cell-autonomously regulates SCPN axon fasciculation within the internal capsule and subsequent pathfinding. Further, integrated investigation of *Bcl11b*-null SCPN with transcriptomic, immunocytochemical, and *in vivo* growth cone purification approaches identifies that Cdh13 is localized along axons and on growth cone surfaces of SCPN *in vivo*, and mediates Bcl11b regulation of SCPN axonal outgrowth. Together, these results demonstrate that Bcl11b controls multiple aspects of SCPN axon development by coordinating intrinsic SCPN cell autonomous subcellular mechanisms and extrinsic MSN non-cell-autonomous mechanisms.

## INTRODUCTION

Axonal connectivity is a critical foundation for CNS function. During CNS development, axons of projection neurons extend from cell bodies (somata), undergo long-distance growth, fasciculate, defasciculate, and collateralize, forming specific circuits in distant target areas.^1^ Projection neuron axons navigate through a constantly changing *in vivo* environment. Correct targeting of axons and their collaterals to form functional circuitry is a complex, multistep, multi-component process integrating often interdependent mechanisms: axon growth; attractive and repulsive contact-mediated and diffusible axon guidance at major and minor trajectory choice points; rate of progression across targets while sensing, testing, and rejecting a myriad of alternative but inappropriate targets; tissue and ECM penetration; and often fasciculation within large axon tracts such as the corpus callosum either with sister axons with the same overall projection trajectory or/and with complementary axons of neurons with the opposite projection trajectory.^2-4^ These steps eventually guide axons to desired targets for further circuit refinement. In the cerebral cortex, axons originating from projection neurons with distinct subtype specificity and areal identity often exhibit distinct responses to the same environment with its mixture of cues regulating growth, attraction, repulsion, and target specificity. Thus, axon guidance is controlled by an interplay between intrinsic neuronal subtype-specific molecular programs, and extrinsic environmental cues.

Corticofugal projection neurons (CFuPN) are a broad class of cortical output neurons that extend their axons away from the cortex, through the internal capsule in the striatum, to targets in subcortical (e.g., thalamus) and subcerebral (e.g., brainstem and spinal cord) regions. Two dominant CFuPN subtypes are corticothalamic projection neurons (CThPN), which reside primarily in layer VI and project axons to specific nuclei of the thalamus, and subcerebral projection neurons (SCPN), which reside in layer V, and extend primary axons to targets caudal to the cerebrum.^5,6^ SCPN include the major and highly clinically relevant subset of corticospinal neurons (CSN).^7-10^ After exiting the cortex, CFuPN axons enter the striatum, fasciculating in the internal capsule via interaction with striatal medium spiny neurons (MSN; also known as striatal projection neurons). Caudal to the striatum, CFuPN axons diverge into thalamic-projecting CThPN axons and subcerebral-projecting SCPN axons projecting to brainstem and/or spinal cord. CThPN axons reciprocally interact with, and are thus guided by, thalamocortical axons to innervate thalamus.^11-13^ A recent study identified that SCPN axons exhibit convergent projection with pioneering MSN axons in the forebrain.^14^ SCPN axons traverse a substantial further distance caudally, forming the corticospinal tract (CST), to reach the brainstem and spinal cord.^15^ Very recent work regarding developing inter-hemispheric callosal axons in the corpus callosum reveals that axonal adhesion molecule levels instruct topographic axon guidance and targeting.^2^ These examples highlight the importance of coordinated regulation between intrinsic and extrinsic molecular elements that together guide axons of distinct neuron populations and subtypes, but little is known about how this molecular coordination is achieved.

The zinc finger transcription factor Bcl11b/Ctip2 is expressed by SCPN at high level postmitotically, and it functions centrally in controlling their differentiation.^16^ Many other molecular-genetic controls over neocortical projection neuron differentiation operate at least in part by regulating Bcl11b expression either positively or negatively.^17-20^ Bcl11b regulates transcription via both DNA binding and nucleosome remodeling.^21,22^ In constitutive *Bcl11b* knockout (*Bcl11b*^-/-^) mice, SCPN are born, express SCPN upstream control genes such as Fezf2, and migrate to layer V, but they exhibit remarkable defects in axon pathfinding, fasciculation, and outgrowth.^16^ SCPN axons normally fasciculate in the internal capsule, the myelinated forebrain tract through the striatum containing many fascicles of axons that project from cortex to subcortical targets. In *Bcl11b*^-/-^ mice, descending axons become misrouted and defasciculated as they pass through the forebrain. While SCPN axons efficiently project to the spinal cord in wild-type mice, SCPN in *Bcl11b*^-/-^ mice extend axons to ectopic targets.^16^ These mutant axons are clearly dysmorphic, and often possess bulbous structures suggestive of dysfunctional growth cones (GCs).^23^ Most critically, SCPN axons in *Bcl11b*^-/-^ mice do not successfully reach the spinal cord; the projections that exit the forebrain mostly terminate in the midbrain, with only rare axons reaching pons, and none reaching the pyramidal decussation.^16^ These findings strongly indicate SCPN-autonomous Bcl11b functions in their axon development.

Intriguingly, Bcl11b is also expressed at high levels by MSN in the striatum, and *Bcl11b*^-/-^ MSN lose their characteristic patch-matrix organization and mis-express key axon guidance molecules.^24^ Because MSN surround SCPN axons as they penetrate through the internal capsule, potentially providing important guidance cues to SCPN, and because the pallial-subpallial boundary (the embryonic border between cortex and striatum) is an important decision point for axons traveling through the internal capsule,^25^ we hypothesized that Bcl11b might also contribute to SCPN development non-SCPN-autonomously via functions in MSN.

While a number of transcriptional controls regulating cortical projection neuron subtype specification and areal development have been identified, downstream subcellular mechanisms and molecular machinery that implement subtype-specific circuitry remain largely unknown. Importantly, previous studies provide evidence that complex subcellular molecular networks are axon GC-localized, including axon guidance ligands, receptors, and adhesion molecules,^1^ and that tightly regulated local translation of select mRNA transcripts directly controls GC pathfinding.^26-28^ Recent *in vivo* work from our lab identifies GC-specific enrichment of hundreds of transcripts and proteins compared with parent somata of developing interhemispheric callosal projection neurons (CPN) and CThPN,^2,29-32^ and demonstrates that subcellular GC molecular machinery directly implements subtype-specific developmental programs to build circuitry, connect to appropriate targets, and select correct synaptic partners. However, subcellular GC molecular machinery underlying SCPN circuit formation remains unknown.

Here, we identify that Bcl11b contributes to SCPN axon development both SCPN-autonomously and SCPN-non-autonomously (via functions in MSN) by employing cortex-specific (*Emx1-Cre*) and/or striatum-specific (*Gsx2-Cre*) *Bcl11b* conditional deletion. The results indicate that Bcl11b orchestrates SCPN axon outgrowth, fasciculation, and pathfinding through complementary functions in two distinct but physically interacting neuron populations (SCPN and MSN). Further, to directly investigate subcellular molecular mechanisms by which Bcl11b cell-intrinsically controls SCPN axon projection targeting, we purified SCPN axonal GCs and somata *in vivo* from developing wild-type and cortex-specific *Bcl11b* knockout mice, and investigated their local subcellular molecular machinery. We identify that Cadherin 13 (Cdh13) is a key downstream target of Bcl11b in SCPN for their axon outgrowth. Cdh13 is expressed by SCPN under control of Bcl11b; is subcellularly localized along SCPN axons and on SCPN GC surfaces *in vivo*; and promotes SCPN axon extension. Together, these results demonstrate that Bcll1b controls multiple aspects of precise SCPN axon development by coordinated intrinsic and extrinsic mechanisms.

## RESULTS

### *Emx1-IRES-Cre* and *Gsx2-iCre* efficiently delete Bcl11b from neocortical neurons and MSN, respectively

In developing forebrain, Bcl11b is expressed at high level by postmitotic SCPN in dorsal cortex, and by MSN in striatum (Figure 1A).^16,24^ To delete Bcl11b specifically from glutamatergic cortical projection neurons, we crossed a *loxP*-flanked (floxed) *Bcl11b* mouse line^33^ with an *Emx1*^*IRES-Cre*^ knock-in mouse line.^34^ These *Bcl11b*^*flox/flox*^*;Emx1*^*IRES-Cre/+*^ mice are born in expected Mendelian ratios, and survive until adulthood, unlike *Bcl11b*^-/-^ mice, which die at postnatal day (P)0.^16,35^ *Bcl11b*^*flox/flox*^*;Emx1*^*IRES-Cre/+*^ tissue does not contain detectable levels of Bcl11b in layer V pyramidal neurons, while Bcl11b expression by MSN is preserved (Figure 1B).

**Figure 1.**
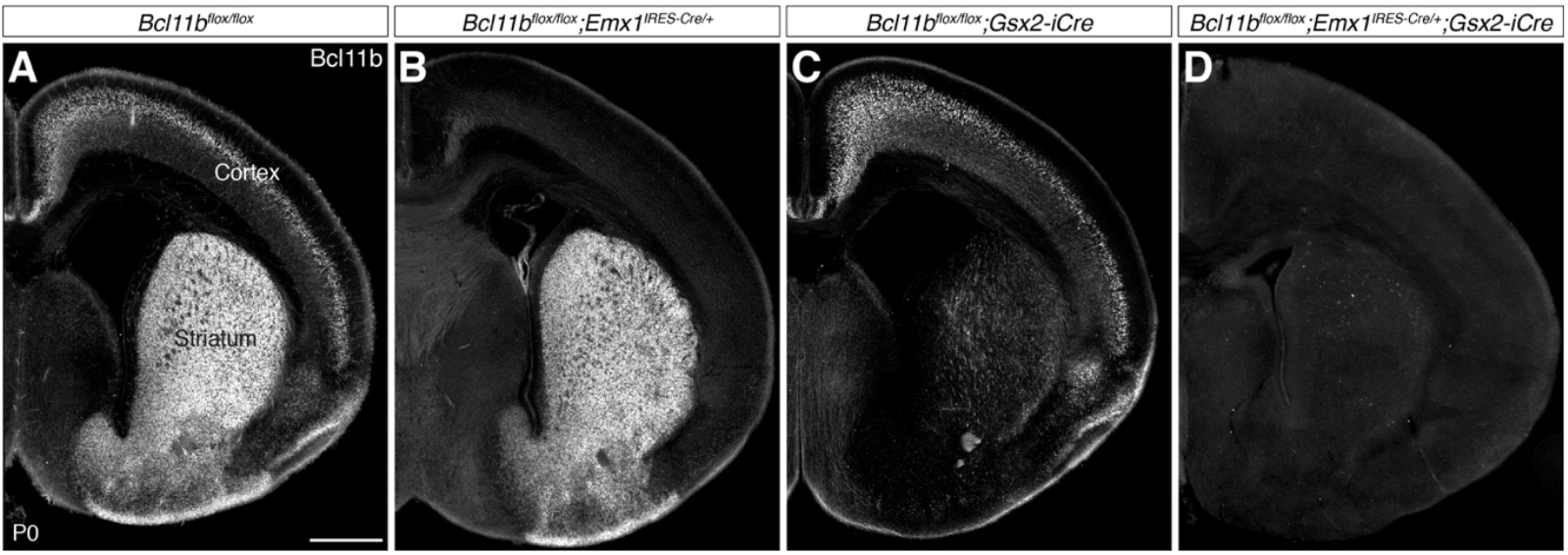
*Emx1-Cre* deletes *Bcl11b* from cortex, and *Gsx2-iCre* deletes *Bcl11b* from striatum. (**A**) Immunocytochemistry on P0 coronal forebrain section detects Bcl11b expression by cortical projection neurons in the cortical plate and by MSN in the striatum of wild-type (*Bcl11b*^*flox/flox*^) mice. (**B**) Bcl11b expression by cortical projection neurons is deleted by *Emx1-Cre*. (**C**) Bcl11b expression by MSN is deleted by *Gsx2-iCre*. (**D**) Double conditional deletion of *Bcl11b* by *Emx1-Cre* and *Gsx2-iCre* deletes Bcl11b from both cortical projection neurons and MSN. Scale bar, 500 µm.

To directly examine MSN-specific Bcl11b functions in SCPN axon development, we crossed the *Bcl11b*^*flox/flox*^ line with a *Gsx2-iCre* transgenic mouse line (*Gsx2* was formerly called *Gsh2*).^36,37^ When *Gsx2-iCre* is crossed with a *Rosa26*^*LSL-LacZ*^ reporter line,^38^ no Bcl11b-positive pyramidal neurons in cortex express β-galactosidase (Supplementary Figure 1A). Interneurons derived from medial ganglionic eminence, identified by immunocytochemistry for parvalbumin, somatostatin, and neuropeptide Y,^39^ are the only β-galactosidase-positive neurons in cortex (Supplementary Figure 1B–D). *Bcl11b*^*flox/flox*^*;Gsx2-iCre* mice are born in expected Mendelian ratios, and survive until adulthood. Bcl11b expression is normal in *Bcl11b*^*flox/flox*^*;Gsx2-iCre* cortex, but is abolished in striatum (Figure 1C). Expression of the MSN marker DARPP-32 is reduced in *Bcl11b*^*flox/flox*^*;Gsx2-iCre* striatum, as reported for *Bcl11b*^*-/-*^ striatum (Supplementary Figure 1E).^24^

When *Bcl11b*^*flox/flox*^ mice are crossed with mice carrying both *Emx1*^*IRES-Cre*^ and *Gsx2-iCre* alleles, the vast majority of Bcl11b expression in the forebrain is abolished (Figure 1D), indicating that Bcl11b is predominantly expressed by *Emx1*-expressing and *Gsx2*-expressing cell lineages. We next used these single- and double-conditional *Bcl11b* mutants to investigate Bcl11b functions in SCPN axon development.

### *Bcl11b* deletion from MSN causes abnormal CFuPN axon fasciculation in the internal capsule

To investigate abnormalities in axon tract morphology and cellular organization within the forebrain due to loss of *Bcl11b* expression by SCPN and/or MSN, we performed Nissl stains (Figure 2A–E) and GAP43 immunocytochemistry (Figure 2F–J; Supplementary Figure 1E) on P0 forebrain tissue sections from wild-type (*Bcl11b*^*flox/flox*^), *Bcl11b*^*flox/flox*^*;Emx1*^*IRES-Cre/+*^, *Bcl11b*^*flox/flox*^*;Gsx2-iCre, Bcl11b*^*flox/flox*^*;Emx1*^*IRES-Cre/+*^*;Gsx2-iCre*, and *Bcl11b*^*-/-*^ mice.

**Figure 2.**
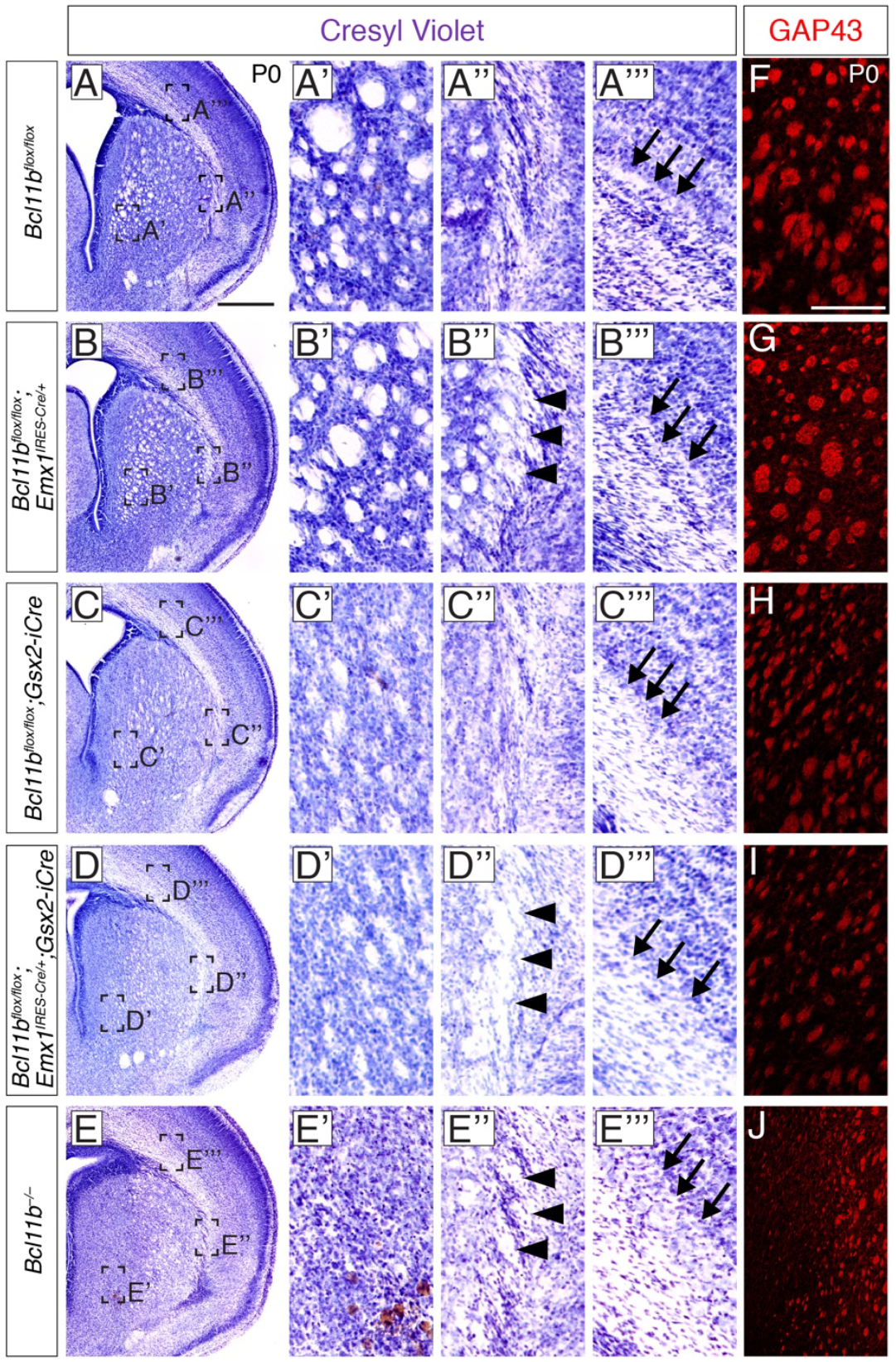
*Bcl11b* expression by MSN in striatum, but not by projection neurons in cortex, is required for CFuPN axon fasciculation within the internal capsule. (**A–A’’’, F**) CFuPN axons exit cortical plate, enter striatum, and fasciculate in the internal capsule in wild-type (*Bcl11b*^*flox/flox*^) mice. (**B, B’, G**) In *Bcl11b*^*flox/flox*^*;Emx1*^*IRES-Cre/+*^ brains, CFuPN axons lacking *Bcl11b* form normal fascicles through striatum, but have an abnormal border between subplate and cortical white matter (B’’’), and an ectopic external capsule (B’’). (**C, C’, H**) In *Bcl11b*^*flox/flox*^*;Gsx2-iCre* brains, CFuPN axon fasciculation is disturbed in striatum, while the border between subplate and cortical white matter remains normal (C’’’), and CFuPN axons enter striatum normally (C’’). (**D, E, I, J**) Brains lacking *Bcl11b* in both cortex and striatum (**D**–**D’’’, E**–**E’’’**) have an abnormal border between subplate and cortical white matter (D’’’, E’’’), an ectopic external capsule (D’’, E’’), and abnormally fasciculated CFuPN axons (D’, E’, I, J). A–E, Nissl staining. Scale bar, 500 µm. F–J, GAP43 immunocytochemistry. Scale bar, 100 µm. Arrowheads indicate ectopic external capsule at the border between cortex and striatum. Arrows indicate location of the border between subplate and cortical white matter.

Cortical projection neurons are located within the cortical plate, and their efferent axons project through the cortical white matter. Wild-type cortex exhibits clear separation of the cortical plate and white matter (Figure 2A’’’). While this cortical structure is unchanged when *Bcl11b* is deleted only from striatum (Figure 2C’’’), the separation becomes less clear when *Bcl11b* is deleted from cortex (Figure 2B’’’, D’’’, E’’’), indicating cell-autonomous *Bcl11b* function in cortical projection neurons for their early axon development within cortex, and for their initial organization toward the internal capsule.

In wild-type mice, CFuPN axons then penetrate into striatum, and fasciculate to form the internal capsule (Figure 2A’, A”, F; Supplementary Figure 1E). When *Bcl11b* is deleted from cortex, those CFuPN axon fascicles that succeed in entering the striatal internal capsule appear normal (Figure 2B’, G; Supplementary Figure 1E), indicating that Bcl11b expression in cortex is largely dispensable for CFuPN axons to fasciculate in striatum. However, organization and entry into the internal capsule is affected: an ectopic external capsule emerges at the lateral border of striatum (Figure 2B”), indicating that cortical *Bcl11b* is necessary for CFuPN axons to optimally penetrate into the striatum. This ectopic external capsule emerges when *Bcl11b* is deleted from cortex (Figure 2B”, D”, E”), but not when it is deleted from striatum (Figure 2C”). Overall, *Bcl11b* expressed in cortex is dispensable for fasciculation of CFuPN axons that successfully enter striatum.

We previously reported that cellular architecture in striatum is substantially disrupted in *Bcl11b*^-/-^ mice.^24^ Indeed, deletion of *Bcl11b* specifically from striatum leads to severely disrupted patch-matrix organization (Figure 2C’–E’). We find disorganized internal capsule structure (Figure 2C’–E’), and impeded CFuPN fasiculation (Figure 2H–J; Supplementary Figure 1E). These results demonstrate that *Bcl11b* expression in striatum is required for CFuPN axon fasciculation in the internal capsule, indicating that *Bcl11b* expressed by MSN importantly regulates CFuPN axon fasciculation non-cell-autonomously.

### *Bcl11b* deletion from SCPN causes reduced SCPN axon outgrowth in brainstem, and *Bcl11b* deletion from MSN causes ectopic entry of SCPN axons into subthalamic nucleus and dorsocaudal midbrain

To investigate how *Bcl11b* deletion from distinct neuronal populations affects SCPN axon development caudal to striatum, we anterogradely labeled SCPN axons using a genetic reporter. We crossed *Bcl11b* wild-type or mutant mice with *Emx1*^*IRES-Flpo*^*;Rosa26*^*frt-LacZ-frt-EGFP*^ mice^8,40^ to broadly label cortical projection neurons by EGFP, independent of Cre expression status (Figure 3A). At P1, EGFP expression is indistinguishable between wild-type and any of the *Bcl11b* mutant cortices (Supplementary Figure 2A–D). EGFP expression is mosaic with a dorsal-to-ventral gradient, reflecting a graded expression pattern of Emx1.^41^ CFuPN axons form a large bundle of axons after exiting striatum, then a subset (CThPN axons) innervate thalamus, while the rest (SCPN axons) extend caudal to thalamus (Supplementary Figure 2E–H). Despite low-level Bcl11b expression by CThPN,^16^ innervation of CThPN axons into thalamus appears essentially normal in all mutant mice compared to wild-type mice (Supplementary Figure 2E–H). We next focused on and deeply investigated SCPN axon trajectory.

**Figure 3.**
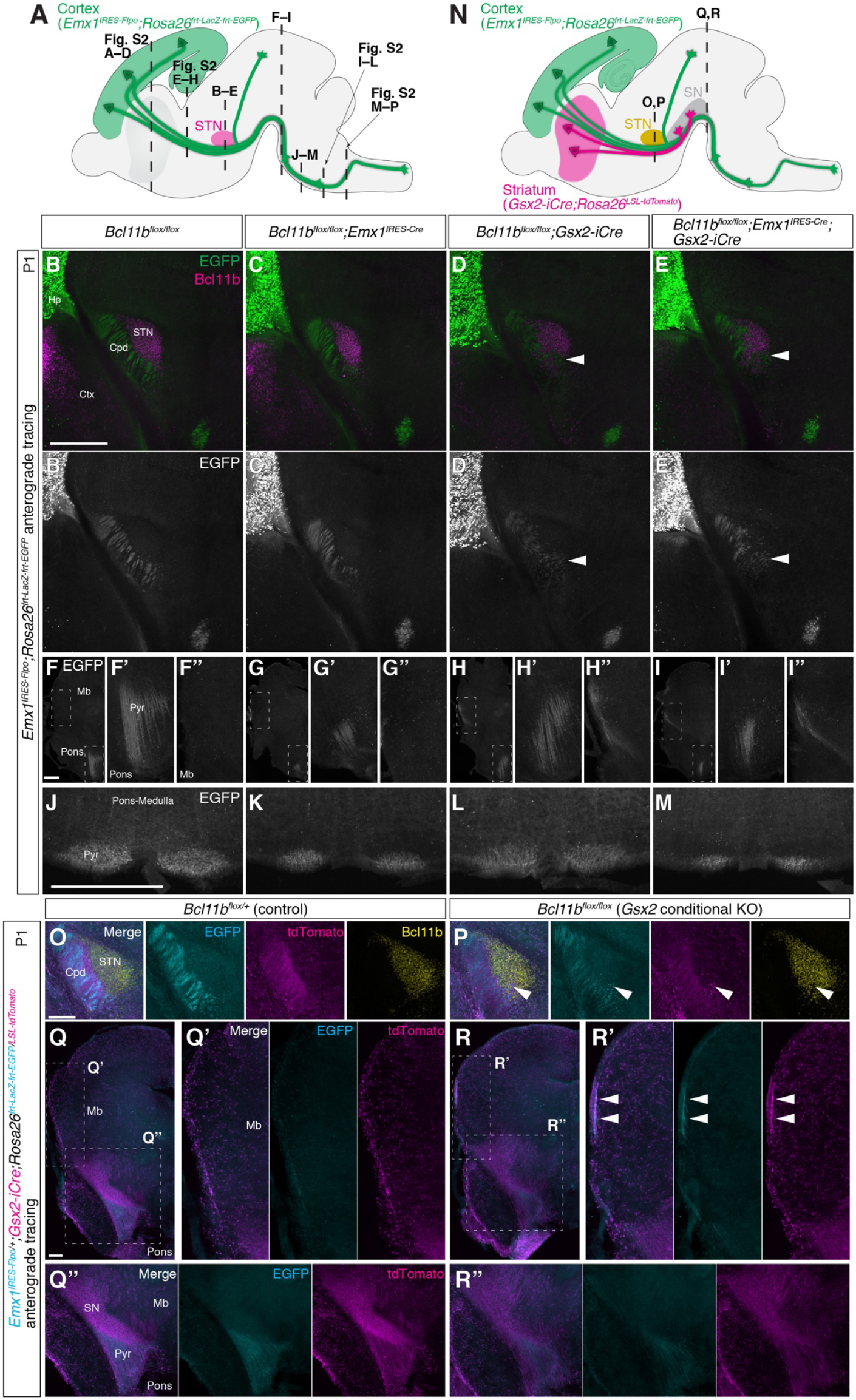
*Bcl11b* deletion from SCPN disrupts outgrowth of SCPN axons, and *Bcl11b* deletion from MSN causes SCPN axon mistargeting into STN and dorsocaudal midbrain. (**A**–**M**) Cortical efferent axons are visualized anterogradely by *Emx1*^*IRES-Flpo*^*;Rosa26*^*frt-LacZ-frt-EGFP*^ in coronal sections of P1 brains. (A) Schematic illustrates positions of coronal sections. (B–M) EGFP-positive SCPN axons (green or white) and Bcl11b (magenta) are shown. While SCPN axons project laterally to STN in *Bcl11b*^*flox/flox*^ and *Bcl11b*^*flox/flox*^*;Emx1*^*IRES-Cre/+*^ brains (B, C), SCPN axons mis-project into Bcl11b-labeled STN when Bcl11b is deleted from striatum (D, E; white arrowheads). (F) In *Bcl11b*^*flox/flox*^ mice, all SCPN axons enter pons (F’) and do not project into the midbrain area dorsal to pons (F”). (G) In *Bcl11b*^*flox/flox*^*;Emx1*^*IRES-Cre/+*^ mice, SCPN axons entering pons are substantially reduced (G’), and do not project into the midbrain area dorsal to pons (G”). (H) In *Bcl11b*^*flox/flox*^*;Gsx2-iCre* mice, SCPN axons abnormally target into midbrain (H”), and there is a slight reduction of SCPN axons entering pons (H’). (I) In *Bcl11b*^*flox/flox*^*;Emx1*^*IRES-Cre/+*^*;Gsx2-iCre* mice, SCPN axons both have reduced entry into pons (I’) and abnormally target into midbrain (I”). Consequently, compared to control (J), SCPN axons reaching medial and caudal brainstem are substantially reduced in *Bcl11b*^*flox/flox*^*;Emx1*^*IRES-Cre/+*^ mice (K), and are slightly reduced in *Bcl11b*^*flox/flox*^*;Gsx2-iCre* mice (L). Deletion of *Bcl11b* from both cortex and striatum additively reduces SCPN axons reaching medial and caudal brainstem (M). See also Supplementary Figure 2A–P. Scale bars, 500 µm. (**N**–**R**) Simultaneous visualization of cortical efferent axons by *Emx1*^*IRES-Flpo*^*;Rosa26*^*frt-LacZ-frt-EGFP*^ and MSN axons by *Gsx2-iCre;Rosa26*^*LSL-tdTomato*^ in coronal sections of P1 brains. (N) Schematic illustrates positions of coronal sections. EGFP, cyan. tdTomato, magenta. Bcl11b, yellow. (O) In control brains, EGFP-positive SCPN axons and tdTomato-positive MSN axons project together lateral to STN, which expresses Bcl11b. (Q, Q’, Q”) These EGFP-positive SCPN axons subsequently project into pons at the junction of midbrain and hindbrain surrounded by tdTomato-positive axons. (P) In Gsx2 conditional KO, a subset of axons project abnormally into STN— all are positive for both EGFP and tdTomato (white arrowheads). (R, R’) EGFP-positive SCPN axons abnormally targeting into dorsocaudal midbrain colocalize with similarly mistargeting tdTomato-positive MSN axons (arrowheads). (R, R”) MSN axons reaching substantia nigra and SCPN axons entering pons are reduced compared to control. Scale bars, 200 µm. Cpd, cerebral peduncle. Ctx, cortex. Hp, hippocampus. STN, subthalamus nucleus. Mb, midbrain. Pyr, pyramidal tract. SN, substantia nigra.

In wild-type mice, SCPN axons fasciculate tightly and form the cerebral peduncle in the midbrain, projecting laterally to the subthalamic nucleus (STN; STN neurons also express Bcl11b), and extending caudally (Figure 3B). These SCPN axons eventually enter the pons (Figure 3F, F’), and no SCPN axons project toward dorsocaudal midbrain (Figure 3F”). SCPN axons continue extending caudally to reach the medulla (Figure 3J; Supplementary Figure 2I); many form the pyramidal decussation (Supplementary Figure 2M).

SCPN axons in *Bcl11b*^*flox/flox*^*;Emx1*^*IRES-Cre/+*^ mice fasciculate normally in the cerebral peduncle and project laterally to STN (Figure 3C). No SCPN axons target dorsocaudal midbrain, but very few SCPN axons reach pons and medulla, resulting in a substantially underdeveloped pyramidal decussation (Figure 3G, G’, G”, K; Supplementary Figure 2J, N; see also Supplementary Figure 5). These results indicate that *Bcl11b* expressed by SCPN is required cell-autonomously for SCPN axon outgrowth in midbrain and hindbrain (and thus further caudally).

We next find that SCPN axon targeting is regulated non-cell-autonomously by Bcl11b expressed by MSN. While SCPN axons in *Bcl11b*^*flox/flox*^*;Gsx2-iCre* mice exhibit fasciculation defects within the internal capsule (Figure 2C, C’, C”), the majority of these axons ultimately project through the cerebral peduncle and pons (Figure 3D, H, L; Supplementary Figure 2K, O). However, a small subset of SCPN axons penetrate through the STN, even though Bcl11b expression in the STN remains intact (Figure 3D). Further, a subset of SCPN axons mistargets dorsocaudal midbrain (Figure 3H”), presumably leading to a corresponding reduction of SCPN axons reaching the pons (Figure 3H’, L; Supplementary Figure 2K, O). Quite intriguingly, SCPN axon mistargeting toward dorsocaudal midbrain is not always symmetric between hemispheres: some mice show unilateral mistargeting (1/5 mice), while others show bilateral (4/5 mice), and the severity of bilateral mistargeting is often variable between hemispheres.

In *Bcl11b*^*flox/flox*^*;Emx1*^*IRES-Cre/+*^*;Gsx2-iCre* mice (Figure 3E, I, M; Supplementary Figure 2L, P), defects in SCPN axon pathfinding and outgrowth recapitulate those observed in *Bcl11b*^*flox/flox*^*;Emx1*^*IRES-Cre/+*^ and *Bcl11b*^*flox/flox*^*;Gsx2-iCre* mice, resulting in even fewer SCPN axons extending in the brainstem (Figure 3M; Supplementary Figure 2L, P). These results further reinforce our findings that Bcl11b expressed by SCPN themselves cell-autonomously controls SCPN axon outgrowth, and that Bcl11b expressed by MSN is non-cell-autonomously required for optimal SCPN axon pathfinding.

We further investigated SCPN axon development using an orthogonal anterograde labeling approach: focal DiI injection. We placed DiI crystals in sensorimotor cortex of fixed P0 brain tissue from wild-type, *Bcl11b*^*flox/flox*^*;Emx1*^*IRES-Cre/+*^, *Bcl11b*^*flox/flox*^*;Gsx2-iCre*, and *Bcl11b*^*flox/flox*^*;Emx1*^*IRES-Cre/+*^*;Gsx2-iCre* mice, then visualized anterogradely spread DiI along axons in sagittal sections. We focused on DiI labeled axons projecting more caudal than thalamus, which are unequivocally SCPN axons. We observed essentially identical defects in SCPN axon development (Supplementary Figure 2Q–T) that are identified by our genetic labeling approach (Figure 3A–L), although mistargeting of SCPN axons into STN was not readily noticeable due to sectioning orientation. We additionally identified that *Bcl11b* expressed by SCPN is required for corticotectal branching of SCPN axons toward the optic tectum in the midbrain, and for fasciculation of SCPN axons in the cerebral peduncle (Supplementary Figure 2Q’–T’). DiI stains cell membranes brightly, enabling identification of this SCPN axon defasciculation in the cerebral peduncle.

We next investigated SCPN mistargeting after *Bcl11b* deletion from MSN. Because MSN axons pioneer SCPN axons,^14^ we hypothesized that *Bcl11b* deletion from MSN might perturb their pioneering functions for SCPN axons (while maintaining normal MSN projections), or might mistarget MSN axons, leading to secondary SCPN mistargeting. To distinguish these possibilities, we simultaneously visualized SCPN axons (*Emx1*^*IRES-Flpo*^*;Rosa26*^*frt-LacZ-frt-EGFP*^) and MSN axons (*Gsx2-iCre;Rosa26*^*LSL-tdTomato*^) (Figure 3N). In control mice (*Bcl11b*^*flox/+*^*;Gsx2-iCre;Emx1*^*IRES-Flpo/+*^*;Rosa26*^*frt-LacZ-frt-EGFP/LSL-tdTomato*^), tdTomato-positive MSN axons and EGFP-positive SCPN axons traverse laterally to STN, and no tdTomato- or EGFP-positive axons penetrate inside STN (Figure 3O). In *Gsx2-iCre* conditional knockout mice (*Bcl11b*^*flox/flox*^*;Gsx2-iCre;Emx1*^*IRES-Flpo/+*^; *Rosa26*^*frt-LacZ-frt-EGFP/LSL-tdTomato*^), a small fraction of axons aberrantly pass through STN— all are positive for both tdTomato and EGFP (Figure 3P). Additionally, bundles of both tdTomato-positive and EGFP-positive axons undergo ectopic dorsal misrouting in midbrain of *Gsx2-iCre* conditional null mice (Figure 3Q, Q’, R, R’), which presumably contributes to fewer MSN axons reaching substantia nigra and fewer SCPN axons entering pons (Figure 3Q”, Q”). These findings indicate that mistargeting of *Bcl11b*-null MSN axons secondarily causes mistargeting of wild-type SCPN axons, while Bcl11b expressed by MSN is dispensable for the interaction between MSN and SCPN axons.

Although ectopic expression of Bcl11b is sufficient to direct some CPN axons into the internal capsule,^42^ and although several molecular controls over neocortical projection neuron development act at least in part by suppressing Bcl11b expression in non-SCPN populations,^17,19,20,43^ it is not known whether *Bcl11b* expression by SCPN is necessary for a subset of SCPN axons (i.e., corticospinal axons) to innervate spinal cord, given the perinatal lethality of *Bcl11b*^*-/-*^ mice.^16,35^ To investigate whether *Bcl11b* expressed in the cortex regulates SCPN axon extension into spinal cord, we retrogradely labeled corticospinal axons from cervical spinal cord at P21 using cholera toxin B subunit (CTB). At P25, substantially fewer somata are labeled by CTB in *Bcl11b*^*flox/flox*^*;Emx1*^*IRES-Cre/+*^ cortex than in wild-type cortex (Supplementary Figure 3), indicating that *Bcl11b* is necessary for optimal corticospinal projection.

Taken together, our results indicate that Bcl11b orchestrates SCPN axon development both cell-autonomously via expression by SCPN, and non-cell-autonomously via expression by MSN.

### Bcl11b controls expression of key axon guidance genes in developing SCPN somata

To further investigate cell-autonomous functions of Bcl11b in SCPN axon growth and circuit development, we investigated *in vivo* SCPN transcriptomic changes induced by loss of *Bcl11b* function. We first employed SCPN somata purified by fluorescence-activated cell sorting (FACS).^16,44^ These investigations identify changes in soma transcriptomes (Figure 4; Supplementary Figure 4), which predominantly reflect dysregulated transcription in the nucleus.

**Figure 4.**
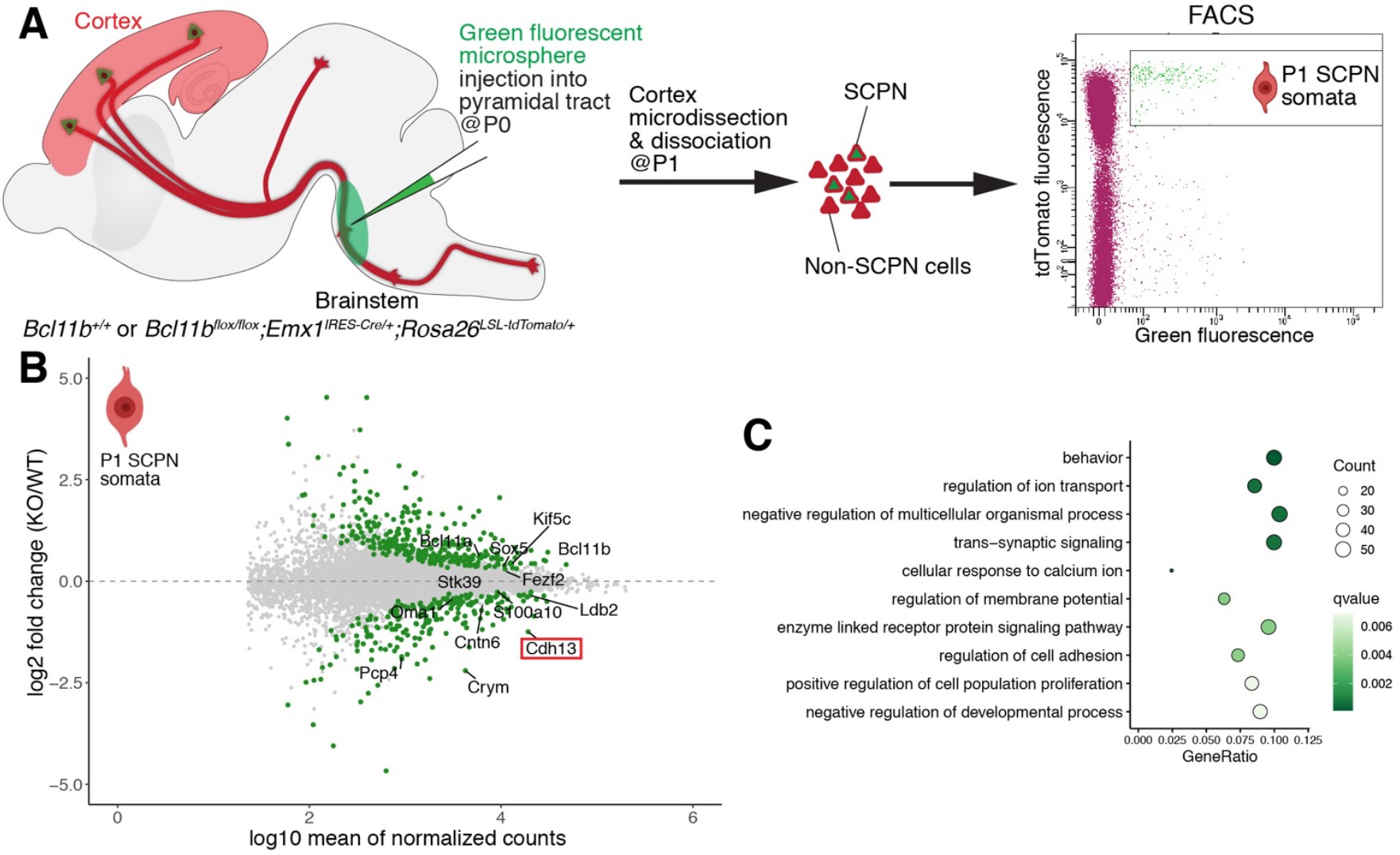
Bcl11b controls expression and subcellular localization of SCPN mRNAs. (A) Schematic for purification of SCPN somata. Green fluorescent microspheres were injected into the pyramidal tract in rostral brainstem of P0 *Emx1-Cre/Rosa26-tdTomat*o pups, and SCPN somata were purified at P1 as cells double positive for both green microspheres and tdTomato. A representative FACS plot is shown. (**B**) Changes to gene expression in SCPN somata upon deletion of *Bcl11b* by *Emx1-Cre*. 515 differentially expressed genes (DEGs; q < 0.1) are shown in green. Other genes are shown in gray. Total: 10,975 genes. SCPN “identity” genes,^16^ *Bcl11a*, and *Kif5c* are labeled. (**C**) GO enrichment analysis of soma DEGs. Points are colored by q-value, and sized by number of genes per term.

To purify SCPN somata, we retrogradely labeled SCPN by injecting green fluorescent microspheres into rostral brainstem of P0 *Bcl11b*^*+/+*^*;Emx1*^*IRES-Cre/+*^*;Rosa26*^*LSL-tdTomato/+*^ and *Bcl11b*^*flox/flox*^*;Emx1*^*IRES-Cre/+*^*;Rosa26*^*LSL-tdTomato/+*^ mice (Supplementary Figure 4A), then isolated cells at P1 that are positive for both tdTomato (i.e., *Emx1*-expressing cortical cells) and green microspheres (i.e., retrogradely labeled neurons; SCPN) via FACS (Figure 4A). Although loss of *Bcl11b* function in the cortex causes substantial reduction of SCPN axons in brainstem (Figure 3; Supplementary Figure 2), the distribution of SCPN retrogradely labeled from rostral brainstem is not markedly different between *Bcl11b*^*flox/+*^*;Emx1*^*IRES-Crel+*^*;Rosa26*^*LSL-tdTomato/+*^ and *Bcl11b*^*flox/flox*^*;Emx1*^*IRES-Crell+*^*;Rosa26*^*LSL-tdTomato/+*^ mice (Supplementary Figure 4B). Retrogradely labeled SCPN in lateral cortex are moderately reduced, but the number of isolated tdTomato/microsphere double-positive neurons per hemisphere is only slightly (but not significantly) reduced by *Bcl11b* deletion from the cortex (Supplementary Figure 4C), indicating that *Bcl11b* is not required for SCPN axon extension into the midbrain/brainstem, despite its necessity for SCPN axon growth and extension after reaching brainstem.

We next performed RNA-seq on P1 FACS-purified SCPN somata to identify potential transcriptomic changes caused by *Bcl11b* deletion from the cortex (Supplementary Tables 1 and 2). Soma transcriptomes cluster by genotype (Supplementary Figure 4D), and demonstrate successful SCPN purification, since wild-type SCPN abundantly express SCPN marker and control genes, including *Fezf2, Cntn6, Cdh13, Sox5, Ldb2, Grb14, Stk39*, and *Crym*.^16^ Interestingly, while exon 2 of the *Bcl11b* gene, which is flanked by *loxP* cassettes, is not detected in SCPN from *Bcl11b*^*flox/flox*^*;Emx1*^*IRES-Cre/+*^ mice, expression of the remaining *Bcl11b* gene is increased by ∼60% in these SCPN (q < 10^-6^) (Supplementary Figure 4E). Since Bcl11b protein is not detected in *Bcl11b*^*flox/flox*^*;Emx1*^*IRES-Cre/+*^ cortex by antibodies against either the N- or C-terminus (Supplementary Figure 4B, F), excision of *Bcl11b* exon 2 efficiently ablates Bcl11b protein expression in the cortex. As expected, *Bcl11a* expression is increased by ∼50% in SCPN from *Bcl11b*^*flox/flox*^*;Emx1*^*IRES-Cre/+*^ mice (q < 10^-6^), since *Bcl11a* and *Bcl11b* are genetically cross-repressive.^18^ The expression of transcriptional regulators with central control over postmitotic cortical projection neuron subtype specification— *Fezf2, Sox5, Tbr1*, and *Satb2*^5^— is indistinguishable between wild-type and *Bcl11b*-null SCPN (Supplementary Table 3), indicating that initial delineation of overall SCPN subtype identity is superficially normal in *Bcl11b*^*flox/flox*^*;Emx1*^*IRES-Cre/+*^ mice. However, expression of other key transcriptional regulators is altered, including the upregulation in *Bcl11b*-null SCPN of CSN marker *Etv1/ER81*^45^ and transcriptional regulators that regulate CThPN differentiation vs. alternate SCPN differentiation— *Zfpm2*/*Fog2* and *Tle4*^46,47^ (Supplementary Table 3). Corticospinal subpopulation-specific genes *Crym* and *Cartpt*^8^ are downregulated by *Bcl11b* deletion, consistent with their functions in corticospinal projection. These dysregulations indicate altered progressive refinement of SCPN subtype identity in the absence of *Bcl11b*.^5^

Overall, expression of 515 genes is significantly altered (q < 0.1) in SCPN by *Bcl11b* deletion from the cortex (Figure 4B; Supplementary Table 2). Of the 515 differentially expressed genes (DEGs), 299 are upregulated and 216 are downregulated (Figure 4B). 35% of previously identified / reported SCPN identity and development genes have significantly altered expression upon *Bcl11b* deletion, while genes known to be expressed by CPN and/or corticotectal neurons, but not by SCPN, remain unchanged (Supplementary Table 4),^16^ supporting Bcl11b’s specific role as a central regulator of SCPN axon development. To identify DEGs known in other systems to be involved in biological processes possibly related to SCPN axon development, we performed gene ontology (GO) term enrichment analysis (Supplementary Table 5). The 10 most significant terms are notably enriched for cell membrane-related biological processes (e.g. trans-synaptic signaling, membrane potential, ion transport, cell adhesion, and receptor protein signaling) (Figure 4C; Supplementary Table 5), indicating that Bcl11b regulates expression of intercellular communication molecules relevant to axon guidance. Therefore, we focused a next stage of investigation on the effect of *Bcl11b* deletion on well-known axon guidance molecules in SCPN. Expression of *Efnb3, Epha4, Ephb1, Ephb6, Plxna1, Sema3C, Draxin*, and *Flrt3* change significantly in *Bcl11b*-null SCPN compared to wild-type SCPN (Supplementary Table 6), suggesting that disrupted transcriptional regulation of axon guidance molecules might contribute importantly to SCPN axon projection defects in *Bcl11b*^*flox/flox*^*;Emx1*^*IRES-Cre/+*^ mice. In addition to these canonical axon guidance molecules, expression of several cadherin genes, including *Cdh13*, reported as an SCPN “general identity” gene,^16^ are differentially expressed by *Bcl11b-*null SCPN (Supplementary Tables 1 and 4).

### Cdh13 is localized to axons and GC surfaces of SCPN *in vivo* in a Bcl11b-dependent manner

Many genes dysregulated with loss of Bcl11b function are involved in intercellular communication, including axon guidance and cell adhesion, but little is known about how these genes, downstream of Bcl11b, function in SCPN axon development. Therefore, we focused investigation on Cdh13 as an exemplar DEG in SCPN axon development.

Cdh13, like other cadherin family members, enables intercellular adhesion via homophilic interactions, but, unlike typical cadherins, it is GPI-anchored, rather than membrane inserted, and does not tether cytoskeletal components or transmit signals on its own.^48,49^ *Cdh13* is highly expressed by wild-type SCPN (233 ± 50 transcripts per million (TPM), Mean ± SD, n = 5), and is reduced by ∼60% in *Bcl11b-*null SCPN (99 ± 20 TPM, n = 5) (Figure 4B). *Cdh13* was previously reported by our lab as a general SCPN “general identity” gene,^16^ meaning in that report strikingly higher level expression by SCPN compared with CPN from early development through postnatal maturation, and suggesting potential importance for establishment and maintenance of identity. However, nothing further is known about its protein expression by SCPN, or its potential function(s) in SCPN.

To investigate Cdh13 protein expression, we immunolabeled Cdh13 at P1. In wild-type cortical plate, Cdh13 is detected along cell membranes, and is more abundant in deep layers than superficial layers (Figure 5A; Supplementary Figure 5). In striatum, Cdh13 signal is widely present and is prominently enriched along axon fascicles in the internal capsule (Figure 5A; Supplementary Figure 5). Cdh13 is abundant along CFuPN axons, including CThPN axons innervating thalamic nuclei, and SCPN axons projecting through the midbrain and hindbrain (Figure 5A, B; Supplementary Figure 5). Notably, Cdh13 immunolabeling along SCPN axon fascicles is stronger in more caudal locations along the corticospinal tract (Figure 5B’; Supplementary Figure 5), suggesting that Cdh13 might be particularly involved in distal SCPN axon growth and extension. In P1 *Bcl11b*^*flox/flox*^*;Emx1*^*IRES-Cre/+*^ mice, Cdh13 immunolabeling in cortex itself is essentially indistinguishable from wild-type (Figure 5C; Supplementary Figure 5), and is unchanged along CFuPN axons in the forebrain, including CThPN axons innervating thalamus (Figure 5C, D; Supplementary Figure 5).

**Figure 5.**
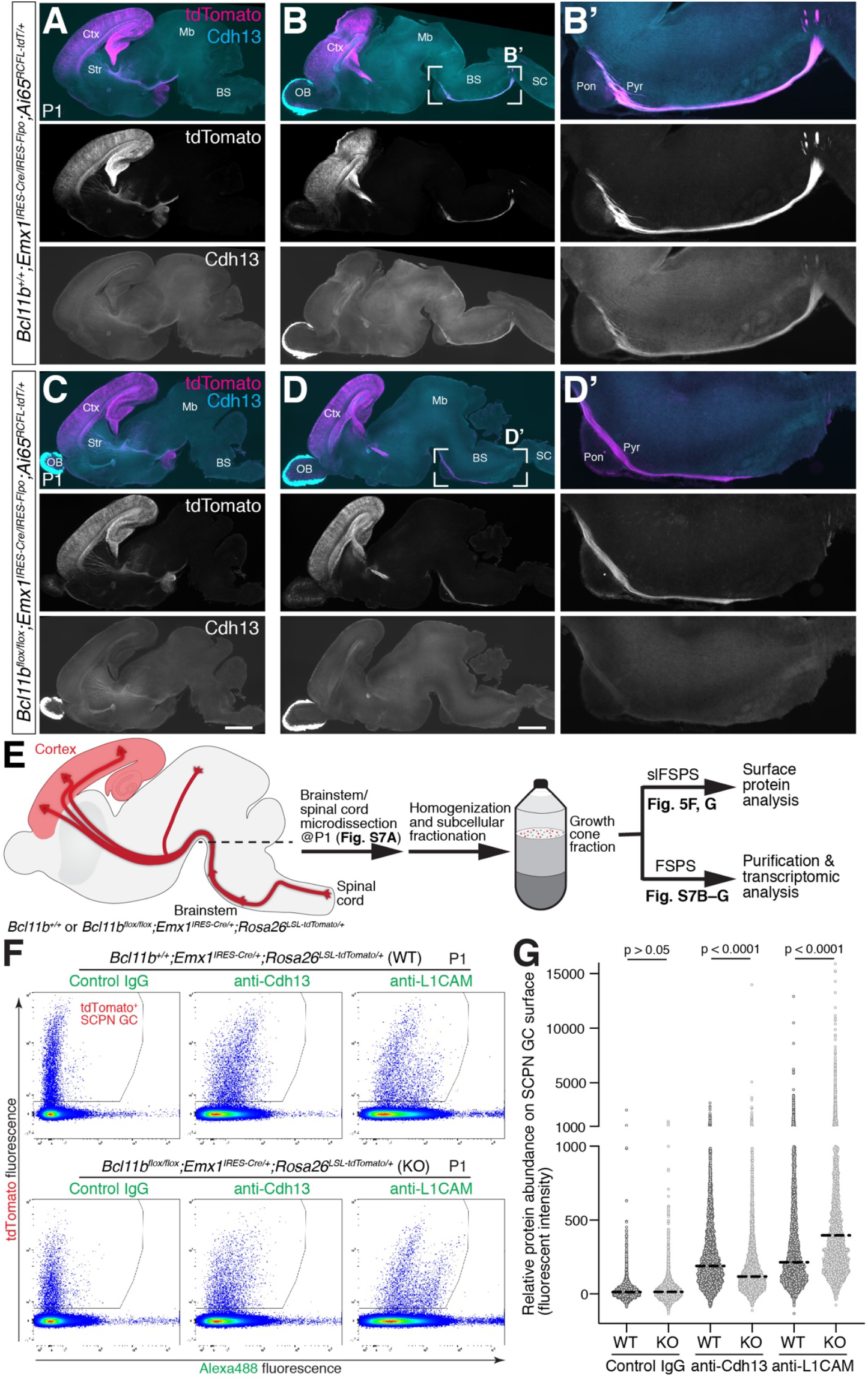
Cdh13 is normally present on SCPN axons and GC surfaces, and *Bcl11b* deletion from SCPN reduces Cdh13 on these subcellular compartments. (**A**–**D**) Immunocytochemistry for Cdh13 (cyan) and tdTomato fluorescence (labeling *Emx-*expressing cell lineage, magenta) in sagittal sections of P1 brains. (A, B, B’) Cdh13 co-localizes with tdTomato-labeled SCPN axons in wild-type brains. (C, D, D’) When *Bcl11b* is deleted from cortex, SCPN axon growth is perturbed, and Cdh13 abundance along SCPN axons is substantially reduced in brainstem. Cdh13 expression in olfactory bulb is unaffected by *Bcl11b* deletion from cortex. Scale bars, 1 mm. (**E**) Schematic for purification of SCPN GCs. Crude growth cone fraction containing SCPN GCs was collected from brainstem and spinal cord of P1 *Emx1-Cre/Rosa26-tdTomat*o pups by subcellular fractionation, followed by fluorescent small particle sorting (FSPS) for surface protein analysis or for purification and transcriptomic investigation of tdTomato-positive SCPN GCs. (**F**) tdTomato-positive SCPN GCs isolated from P1 *Bcl11b*^*+/+*^*;Emx1*^*IRES-Cre/+*^*;Rosa26*^*LSL-tdTomato/+*^ or *Bcl11b*^*flox/flox*^*;Emx1*^*IRES-Cre/+*^*;Rosa26*^*LSL-tdTomato/+*^ brainstem/spinal cord were incubated with control IgG antibody, anti-Cdh13 antibody, or anti-L1CAM antibody, then with Alexa488-conjugated secondary antibody, followed by FSPS. Identical gates are applied for analyzing tdTomato-positive SCPN GCs in (G). (**G**) Quantification was performed for relative protein abundance on tdTomato-positive SCPN GC surfaces *in vivo*. Dotted line indicates median. Mann-Whitney test. BS, brainstem. Ctx, cortex. Str, striatum. Mb, midbrain. OB, olfactory bulb. Pyr, pyramidal tract. SC, spinal cord.

Strikingly, however, Cdh13 immunolabeling along SCPN axons in the brainstem is substantially reduced in *Bcl11b*^*flox/flox*^*;Emx1*^*IRES-Cre/+*^ mice compared to wild-type mice (Figure 5D’; Supplementary Figure 5). To control for the reduction in the number of SCPN axons reaching the brainstem in *Bcl11b*^*flox/flox*^*;Emx1*^*IRES-Cre/+*^ mice, we immunolabeled in parallel for L1CAM, which is not transcriptionally regulated by Bcl11b in SCPN (Supplementary Table 1). Importantly, while L1CAM reduces in proportion to the reduction of SCPN axons in *Bcl11b*^*flox/flox*^*;Emx1*^*IRES-Cre/+*^ mice (Supplementary Figure 5), there is a substantially disproportional decrease of Cdh13. These results indicate that Cdh13 is expressed abundantly by SCPN under the control of Bcl11b, and is localized along SCPN axons *in vivo*, particularly in distal regions of the corticospinal tract.

To investigate whether Cdh13 is present in SCPN GCs *in vivo*, and whether its GC abundance is altered by *Bcl11b* deletion from SCPN, we developed a subtype-specific “surface labeling Fluorescent Small Particle Sorting” (slFSPS) approach that enables quantitative identification of surface antigens on subtype-specific GC membranes *in vivo* (Figure 5E; Supplementary Figure 6A), similar to approaches combining non-specific synaptosome immunolabeling and flow cytometry.^50,51^ In this slFSPS approach, bulk GCs (“GC fraction”, isolated via subcellular fractionation^52^) are incubated with a surface antigen primary antibody, then a fluorescent secondary antibody is added prior to analysis via FSPS (see Materials and Methods).^2,30^ To determine the feasibility of this approach, we overexpressed GPI-anchored EGFP or cytosolic EGFP in CPN by unilateral *in utero* electroporation,^53^ isolated the GC fraction from contralateral hemispheres,^30,31,52^ then incubated this fraction with an anti-GFP antibody. EGFP-GPI GCs are successfully labeled by anti-GFP antibody, and anti-GFP signal is proportional to EGFP fluorescence (Supplementary Figure 6B), indicating that slFSPS is quantitative. Importantly, cytosolic EGFP GCs are not labeled by anti-GFP antibody (Supplementary Figure 6C), demonstrating that slFSPS detects only surface antigens. Additionally, CPN GCs are labeled by anti-NCAM1 antibody (Supplementary Figure 6B, C), consistent with previous work.^2,54^ Thus, slFSPS enables quantitative investigation of subtype-specific GC-surface antigens *in vivo*.

Using slFSPS, we investigated potential presence and abundance of Cdh13 and L1CAM on SCPN GC surfaces in control *Bcl11b*^*+/+*^*;Emx1*^*IRES-Cre/+*^*;Rosa26*^*LSL-tdTomato/+*^ and loss-of-function *Bcl11b*^*flox/flox*^*;Emx1*^*IRES-Cre/+*^*;Rosa26*^*LSL-tdTomato/+*^ mice (Figure 5E–G). We isolated bulk GCs, including fluorescent SCPN GCs, from P1 brainstem and spinal cord. In *Bcl11b*^*+/+*^*;Emx1*^*IRES-Cre/+*^ mice, antibodies against Cdh13 and L1CAM both produce higher fluorescence intensities relative to a control IgG antibody, demonstrating that Cdh13 and L1CAM are present on SCPN GC surfaces. Importantly, Cdh13 abundance on SCPN GC surfaces is reduced in *Bcl11b*^*flox/flox*^*;Emx1*^*IRES-Cre/+*^ mice, while L1CAM abundance is unexpectedly increased on SCPN GC surfaces in *Bcl11b*^*flox/flox*^*;Emx1*^*IRES-Cre/+*^ mice. These findings demonstrate that Cdh13 is both present on normal SCPN GC surfaces *in vivo*, and locally reduced on SCPN GC surfaces upon *Bcl11b* deletion from SCPN.

Nuclear gene expression sets the foundation for cellular specification and function. Importantly, however, cells also transport transcripts and/or proteins to peripheral cellular compartments to tightly control local subcellular functions,^31,55-61^ enabling prompt and targeted local responses to dynamic environments, and liberating distant subcellular compartments from reliance on transcriptional responses in the nucleus.

We therefore investigated potential dysregulation of subcellular GC RNA localization in *Bcl11b-*null SCPN, and asked whether subcellular localization of *Cdh13* RNA is regulated by *Bcl11b*. To investigate GC transcriptomes, we purified fluorescent SCPN GCs at P1 from brainstem and spinal cord of *Bcl11b wild-type;Emx1*^*IRES-Cre/+*^*;Rosa26*^*LSL-tdTomato/+*^ or *Bcl11b*^*flox/flox*^*;Emx1*^*IRES-Cre/+*^*;Rosa26*^*LSL-tdTomato/+*^ mice via established methods^29-32^ for subcellular fractionation and FSPS (Figure 5E; Supplementary Figure 7A, B). Following GC purification, sequencing, and assessment by rigorous quality control metrics (Supplementary Figure 7C–F; Supplementary Tables 7–9; see Materials and Methods),^31,62^ we identified a focused set of 88 genes as “differentially abundant genes” (DAGs; agnostic regarding potential mechanisms (e.g. expression, trafficking, stability etc.)) between *Bcl11b* wild-type and null GCs (q < 0.1; Supplementary Figure 7F, G; Supplementary Tables 8 and 10), among which 46 are upregulated and 42 are downregulated. GO term enrichment analysis of DAGs does not identify overrepresentation of any specific biological process across the sets, likely due to their small size and the particular of GO term statistical analysis. These results indicate that *Bcl11b* is required for appropriate localization of specific mRNAs to SCPN axonal GCs during development. It remains unknown to what extent Bcl11b control of Cdh13 abundance on SCPN GC surfaces is mediated by nuclear *Cdh13* transcription vs. subcellular *Cdh13* transcript localization and translation.

Together, these findings demonstrate that Cdh13 is localized along SCPN axons, with pronounced distal enrichment along brainstem and spinal cord corticospinal tract axons, and is extracellularly presented on GC surfaces of growing SCPN axons in a Bcl11b-dependent manner.

### Cdh13 is a critical downstream target of Bcl11b in SCPN axon extension control

Cdh13 has been implicated in axonal development of the serotonergic system and is required for retinotectal synapse maintenance,^63,64^ but its function in cortical development has not been investigated. To determine whether Cdh13 regulates SCPN axon development, we examined the effect of *Cdh13* suppression on SCPN axon development. We designed 4 guide RNAs (gRNAs) targeting the *Cdh13* transcription start site, and used CRISPR activation (CRISPRa) to screen functional gRNAs.^65^ All 4 gRNAs induce Cdh13 expression (Supplementary Figure 8A). We then selected the most efficient *Cdh13* gRNAs (#2 and #3) for further testing with CRISPR interference (CRISPRi) to inhibit Cdh13 expression.^66^ This two-step screening identified that gRNA #3 efficiently knocks down Cdh13 expression *in vivo* (Supplementary Figure 8B, B’). *In utero* electroporation of control gRNA^67^ at embryonic day (E) 12.5 leads to efficient SCPN axon extension in the cerebral peduncle and brainstem at E18.5 (Figure 6A).

**Figure 6.**
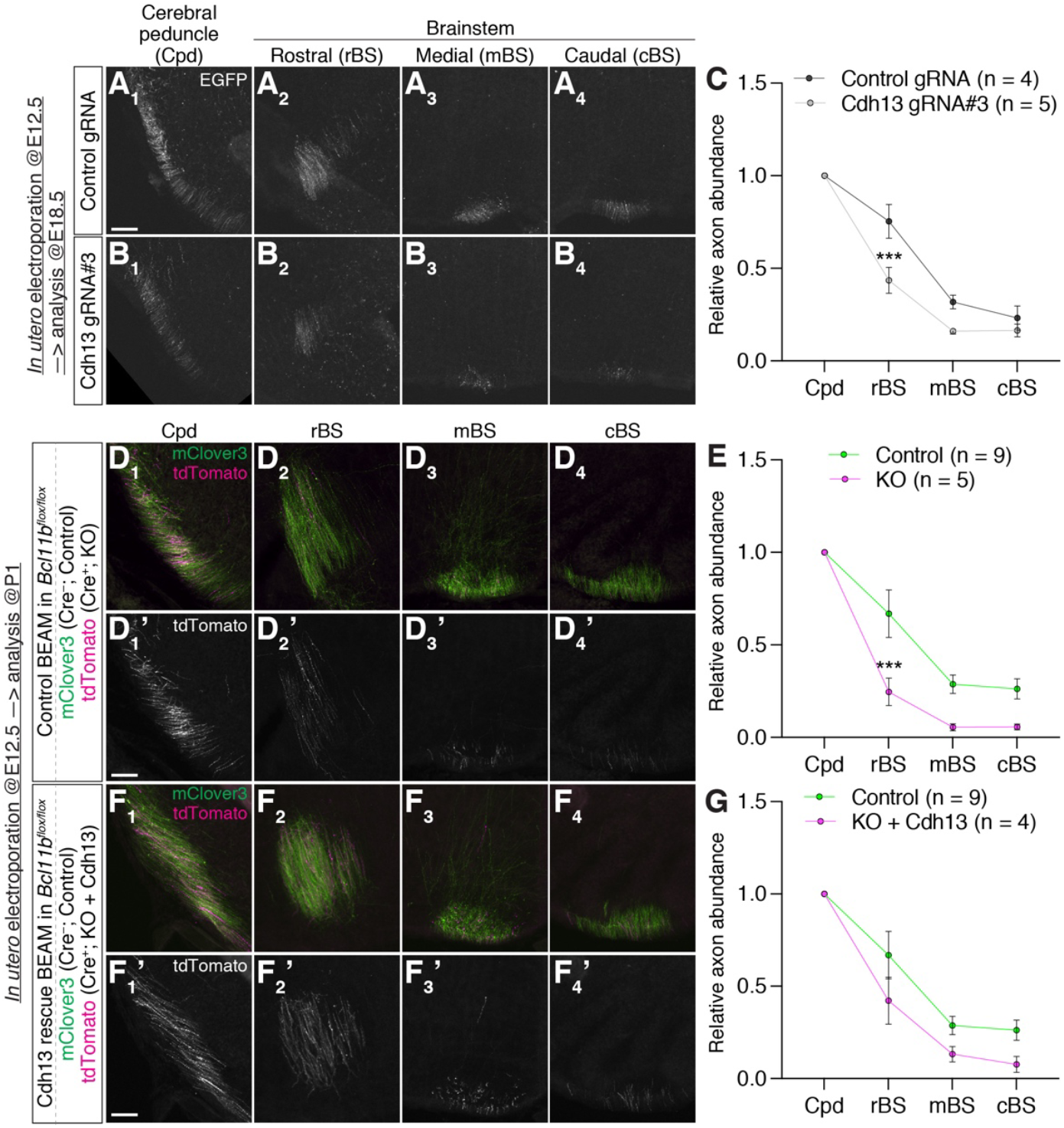
Cdh13 is required for SCPN axon extension in the brainstem, and Cdh13 over-expression ameliorates SCPN axon extension defects induced by cell-autonomous *Bcl11b* deletion. (**A**–**C**) Fluorescence images of E18.5 wild-type brains electroporated *in utero* at E12.5 with a plasmid encoding control gRNA or *Cdh13* gRNA #3, together with those encoding EGFP and dCas9-KRAB-MeCP2. (A, B) EGFP-labeled axons in the cerebral peduncle and brainstem. (C) Quantification at E18.5 of axon abundance in the brainstem relative to axon abundance in the cerebral peduncle reveals that *Cdh13* knockdown disrupts SCPN axon extension in the brainstem. Mean ± SEM. Control gRNA vs. *Cdh13* gRNA #3; p < 0.05, repeated measures two-way ANOVA. ***; p < 0.001, Bonferroni’s multiple comparison test. (**D, E**) Fluorescence images of P1 *Bcl11b*^*flox/flox*^ mouse brains electroporated *in utero* at E12.5 with a precisely-defined cocktail of Cre-recombinase, Cre-inducible Cre-recombinase, Cre-silenced myr-Clover3 (Control; green), and Cre-activated myr-tdTomato (KO; magenta). (D) Fluorescently labeled axons in the cerebral peduncle and brainstem. (E) Quantification of axon abundance in the brainstem relative to axon abundance in the cerebral peduncle reveals that *Bcl11b* knockout disrupts SCPN axon extension in the brainstem. Mean ± SEM. Control vs. KO; p < 0.05, repeated measures two-way ANOVA. ***; p < 0.001, Bonferroni’s multiple comparison test. (**F, G**) Fluorescence images of P1 *Bcl11b*^*flox/flox*^ mouse brains electroporated *in utero* at E12.5 with a precisely-defined cocktail of Cre-recombinase, Cre-inducible Cre-recombinase, Cre-silenced myr-Clover3 (Control; green), Cre-activated myr-tdTomato, and Cre-activated Cdh13 (KO + Cdh13; magenta). (F) Fluorescently labeled axons in the cerebral peduncle and brainstem. (G) Quantification at P1 of axon abundance in the brainstem relative to axon abundance in the cerebral peduncle reveals that Cdh13 over-expression ameliorates SCPN axon extension defects in the brainstem caused by *Bcl11b* knockout. Mean ± SEM. Control vs. KO + Cdh13; p > 0.05, repeated measures two-way ANOVA. Control samples from control BEAM and Cdh13 rescue BEAM experiments are indistinguishable, so they are combined to increase statistical power. Scale bars, 100 µm.

In contrast, reducing *Cdh13* expression by SCPN via gRNA #3 causes a reduced number of axons reaching the cerebral peduncle and brainstem (Figure 6B). Normalizing axon abundance to the axon abundance in the cerebral peduncle demonstrates that *Cdh13* knockdown significantly reduces SCPN axon extension caudal to the cerebral peduncle (Figure 6C). This indicates that Cdh13 importantly regulates SCPN axon extension in the brainstem.

To further investigate in a complementary manner whether *Cdh13* downregulation by the loss of *Bcl11b* in SCPN underlies deficient SCPN axon extension, we overexpressed Cdh13 by *Bcl11b-*null SCPN (Figure 6D–G; Supplementary Figure 8C–E). We performed *in utero* electroporation at E12.5 in *Bcl11b*^*flox/flox*^ mice, using a recently developed Cre recombinase-based mosaic gene manipulation system (Binary Expression Aleatory Mosaic; BEAM),^68^ to simultaneously produce interspersed green-fluorescent, Cre-negative control neurons and red-fluorescent, Cre-positive *Bcl11b* null neurons in the same region of the same cortex (Supplementary Figure 8D). In control BEAM (without Cdh13 overexpression), Cre-negative control SCPN efficiently extend axons into brainstem at P1 (Figure 6D, E). In striking contrast, while Cre-positive SCPN are present in forebrain (Supplementary Figure 8D), their axons are substantially reduced in brainstem (Figure 6D, D’, E), recapitulating the SCPN axon extension defects in *Bcl11b*^*flox/flox*^*;Emx1*^*IRES-Cre/+*^ mice (Figure 3; Supplementary Figure 2). Strikingly, in this Cdh13 rescue experiment employing BEAM, overexpression of Cdh13 only by Cre-positive SCPN partially rescues extension of their axons relative to Cre-negative SCPN (Figure 6F, F’, G; Supplementary Figure 8C– E). These results demonstrate that SCPN axon extension defects caused by the loss of *Bcl11b* in SCPN is mediated, at least in part, by downregulation of Cdh13.

## DISCUSSION

The CST is the longest axon tract in the mammalian CNS, and SCPN axons, including corticospinal axons, execute a particularly complex trajectory through multiple brain regions to reach their final targets in the spinal cord.^15^ SCPN and other CFuPN axons first extend while ventrally repulsed from cortex via Semaphorin signaling,^69,70^ and navigate the pallial-subpallial boundary.^25,37^ Once they enter the internal capsule, CFuPN axons fasciculate with each other, thought to be important for further pathfinding,^15^ then pass the telencephalic-diencephalic boundary.^71^ SCPN axons continue traveling through the midbrain and hindbrain, finishing as axons of CSN reach the pyramidal decussation, cross to the contralateral side, and enter the spinal cord.^72^ The peak birthdate of SCPN is E12.5–13.5 in mouse, and SCPN axons spend the next week alternately growing and pausing until they reach the pyramidal decussation at ∼P0.^73-75^ Because SCPN must interpret a wide range of extracellular signals to arrive at their brainstem and spinal cord targets, investigating interactions between SCPN-autonomous and SCPN-non-autonomous sources of guidance is critical to deepen understanding of SCPN axon development, and potentially to enable directed regeneration.

In this work, we demonstrate that Bcl11b/Ctip2 is not only a critical cell-autonomous regulator of SCPN axon development, but also a substantial and critical non-SCPN-autonomous regulator of SCPN axon projection and tract fasciculation via Bcl11b-expressing MSN, outside the cortex. Within SCPN, Bcl11b is necessary for efficient SCPN axon entry into the internal capsule, corticotectal branching past the internal capsule, and further outgrowth caudal to the midbrain. In contrast to Bcl11b-null mice, which have no SCPN axons reaching the spinal cord (though this absolute lack might be partially due to axon growth delay rather than total disruption of SCPN axon development, since Bcl11b-null mice die soon after birth), deleting *Bcl11b* from only cortex, with normal Bcl11b expression by MSN (*Bcl11b*^*flox/flox*^*;Emx1*^*IRES-Cre/+*^ mice), results in substantial but partial reduction of SCPN axons reaching the spinal cord at P25. Strikingly, deleting *Bcl11b* from only MSN, with normal Bcl11b expression by SCPN (*Bcl11b*^*flox/flox*^*;Gsx2-iCre* mice), strikingly impairs SCPN fasciculation in the internal capsule, and results in other pathfinding defects. Additionally, *Bcl11b* deletion from MSN using *Gsx2-iCre* disorganizes striatal patch-matrix structure,^24^ and alters striato-nigral MSN projection (Figure 3). This is intriguing, since SCPN axons and striato-nigral MSN axons converge and course alongside each other.^14^ Our results indicate that Bcl11b expressed by MSN regulates broad CFuPN axon fasciculation in striatum and continuing SCPN axon pathfinding in midbrain, primarily via control over multi-circuit MSN cellular functions, rather than by only affecting interactions between MSN and CFuPN/SCPN.

Bcl11b regulates transcription via both direct DNA binding and indirect chromatin remodeling, and modulates distinct sets of genes in distinct cell types and contexts.^21,22,76^ To better understand how Bcl11b controls SCPN axon development cell-autonomously, we investigated transcriptomic changes caused by *Bcl11b* deletion from SCPN. We focused on Cdh13, an exemplar differentially expressed gene (DEG) that is continuously expressed by SCPN from early development into maturation^16^ and that has unique molecular features as an atypical cadherin.^48,49,77^ Cdh13 expression and localization at P1 marks the CST. Cdh13 is more enriched in distal portions of SCPN axons. Now, our slFSPS subtype-specific subcellular approaches provide strong evidence that Cdh13 is present directly on SCPN GC surfaces.

Since Cdh13, which is GPI-anchored, is not expected to transmit signals intracellularly on its own, we speculate that Cdh13 serves as a tether to tightly fasciculate SCPN axons via physical forces, thereby enabling long-distance projections of SCPN axons. It is also possible that yet-to-be-discovered co-receptor(s) mediate Cdh13 functions.^78^ Notably, the GPI-anchor biosynthesis gene *Pigq* is a differentially abundant gene (DAG) in null vs. wt SCPN GCs (Supplementary Figure 7G). Importantly, these data reveal that Cdh13 overexpression ameliorates SCPN axon extension defects by *Bcl11b* deletion. Since there might be parallels to the extremely limited axon regrowth of mature CSN/SCPN after CST injury, it would be interesting to investigate whether Cdh13 manipulation might partially promote axon regeneration. Further, *Cdh13* has been implicated in multiple neurodevelopmental disorders, including attention-deficit/hyperactivity disorder and autism spectrum disorder,^79-81^ so it would be interesting to investigate whether altered connectivity of cortical projection neurons contributes to pathogenesis and pathophysiology of such disorders.

The distinct contributions to SCPN axon development of Bcl11b expressed by multiple distinct but physically interacting neuron types highlight both the complexity and the precision of cortical output circuitry evolution. Throughout vertebrate evolution, the CNS has expanded and added more structural complexity rostrally than caudally. SCPN somata are located in cerebral cortex, an advanced brain system specific to mammals, and the CST is a notable feature of all mammals. Only minor projections from the telencephalon to the spinal cord are found in a limited number of non-mammalian vertebrates, including some sharks and birds,^82^ and avian pallium has a small population of neurons with characteristics similar to SCPN, such as expression of SCPN genes (including *Fezf2* and *Bcl11b*), and descending axon projections.^83^ Similar to its expression pattern in mammals, Bcl11b is expressed abundantly in chick striatum.^83^ We speculate that the functions of Bcl11b we identify across distinct-but-interacting neuron types might have initially emerged in a common ancestor of birds and mammals, then evolutionarily enabled in mammals increasingly complex and interactive motor-sensory circuitry. In particular, we speculate that expanded use of these neuronal circuit functions in mammals, in large part via an expanded SCPN population in an enlarged cortex, has contributed substantially to the repertoire of skilled and dexterous voluntary movement in mammals. Interestingly, Bcl11b is also expressed in the subthalamic nucleus (STN), a structure located at the junction of the diencephalon and midbrain, and SCPN axons normally project along the STN border together with the striato-nigral MSN projection.^14^ This suggests that Bcl11b expression in other brain structures might also contribute to guidance of SCPN axons. This raises the more general speculation that single transcriptional regulators beyond Bcl11b might function centrally in distinct cellular populations to enable precise targeting and guidance of long-distance axon projections, thus circuit formation, function, precision, and complexity in the evolution of cortical output circuitry.

How function-specific cortical circuitry is regulated with specificity and precision beyond transcriptional controls is still an underexplored question. This is in large part due to prior inaccessibility of subtype- and stage-specific *in vivo* subcellular RNA and protein molecular machinery. The paired purifications of somata and GCs by subtype-specific FACS and FSPS fills this important gap. As an exemplar of likely combinatorial and multi-element mechanisms, we identify decreased Cdh13 protein on SCPN GC surfaces upon deletion of *Bcl11b* from SCPN (Figure 5F, G). We also unexpectedly identify increased L1CAM protein on *Bcl11b*-null SCPN GC surfaces (Figure 5F, G), even though *L1CAM* transcript abundance in both soma and GC subcellular compartments are unchanged after *Bcl11b* deletion (Supplementary Tables 1 and 8), suggesting that L1CAM protein trafficking to GCs is increased by *Bcl11b* deletion, potentially by altered protein trafficking machinery (see below) and/or by L1CAM substituting in trafficking pathways normally employed for Cdh13.

Further, we identify that deletion of *Bcl11b* causes disruption of subtype-specific axon GC transcriptomes (Supplementary Figure 7), with changes in developmentally and subcellularly functional transcripts likely contributing to disruption of SCPN axonal connectivity, targeting, and organization. Importantly, there is little overlap between DEGs and DAGs (Supplementary Tables 2 and 10). Changes in GC transcriptomes are dependent at the individual transcript level on combinations of not only transcriptional regulation but also or alternatively regulation of subcellular trafficking— GC transcriptomes are often not a direct consequence of nuclear gene expression changes.^29^

Multiple potential molecular mechanisms might underlie altered subcellular transcript and protein abundance in axon GCs of *Bcl11b-*null SCPN. Notably, *Bcl11b* deletion upregulates *Kif5c* expression in SCPN somata (Figure 4B; Supplementary Table 2). Kif5 motor proteins regulate anterograde transport of RNAs, proteins, and organelles,^61,84^ and the motor domain of Kif5c selectively translocates to axonal GCs,^85^ indicating that local molecular machinery in *Bcl11b-*null SCPN GCs might be selectively altered, at least in part, by changes in selective anterograde trafficking. Dysregulation of RNA binding proteins (RBPs) that are critical for transcript-dependent RNA stability and trafficking to GCs might also play a key role — nine RBPs are differentially expressed in *Bcl11b-*null SCPN somata (*Ncl, Csdc2, Zc3h13, Ralyl, Rbm24, Pcbp2, Rbms1, Raver2, Larp1b*), while no RBP RNAs are differentially abundant in GCs, consistent with somatic translation of most RBPs. We did not identify enrichment of 3’ UTR motifs that are bound by RBPs, presumably due to both the small list of DAGs and the currently limited sets of RBP motifs and/or experimentally validated RBP binding sites reported in the literature. Further investigation offers substantial promise to elucidate mechanisms by which Bcl11b controls corticospinal GC subcellular transcriptomes and proteomes, thus corticospinal circuit development and function, and potentially to more broadly elucidate generalizable mechanisms of subtype- and function-specific circuit development, function, dysfunction, disease, and approaches to regeneration.

## SUPPLEMENTAL INFORMATION

Supplementary Figures 1–8

Supplementary Tables 1–10

## ACKNOWLEDGEMENTS

We thank J. Heo, K. Liu, H. McKee, M. Ross, B. Wall, and G. Wheeler for superb technical assistance; K. Ozkan for schematics; members of the Macklis laboratory for scientific discussions and helpful suggestions; N. Kheradmand and J. LaVecchio of the HSCRB-HSCI Flow Cytometry core; the Harvard Bauer RNA sequencing core; the Harvard Center for Biological Imaging for infrastructure and support; R. Kominami, Y. Katsuragi, N. Kessaris, and D. Price for generous sharing of mice. This work was supported by grants from the National Institutes of Health (R01s NS045523 and NS075672, with additional infrastructure supported by DP1 NS106665, NS049553, and NS104055), the Max and Anne Wien Professor of Life Sciences fund, an Allen Distinguished Investigator award from the Paul G. Allen Frontiers Group, and the Massachusetts Department of Public Health Spinal Cord Injury Fund to J.D.M.. Y.I. was partially supported by fellowships from the Uehara Memorial Foundation, Kanae Foundation for the Promotion of Medical Science, the Murata Overseas Scholarship Foundation, and by the DEARS Foundation (to J.D.M.). M.B.W. was partially supported by NIH individual predoctoral National Research Service Award NS064730 and the DEARS Foundation. L.C.G. was partially supported by the Harvard Medical Scientist Training Program, NIH individual predoctoral National Research Service Award NS080343, and the DEARS Foundation. A.K.E. was supported by a Swiss National Science Foundation Postdoctoral Fellowship and a Jean-Jacques et Felicia Lopez-Loreta Foundation Award. D.T. was partially supported by a 2022 NSF-Simon Center at Harvard University QBio Graduate Student Award and NIH NRSA F31 NS127518. J.J.H. was partially supported by NIH NRSA F31 NS103262 and NIH Training Grant T32 GM007226.

## AUTHOR CONTRIBUTIONS

Research designed by Y.I., M.B.W., L.C.G., and J.D.M.; research performed by Y.I., M.B.W., L.C.G., A.K.E., D.T., and J.J.H.; data analyzed by Y.I., M.B.W., L.C.G., D.T., and J.D.M.; and manuscript written by Y.I., M.B.W., and J.D.M.

## EXPERIMENTAL PROCEDURES

### Mice

All mouse work was approved by the Harvard University Institutional Animal Care and Use Committee, and was performed in accordance with institutional and federal guidelines. The date of vaginal plug detection was designated E0.5, and the day of birth as P0. The genders of early postnatal mice were not determined. Mice were group housed with light on a 12:12 or 14:10 h cycle. Water and food were provided *ad libitum*.

Wild-type mice on a C57BL/6J or CD1 background were obtained from Charles River Laboratories (Wilmington, MA). *Bcl11b*^*-/-*^ and *Bcl11b*^*flox/flox*^ mice were the generous gift of Prof. Ryo Kominami.^33,35^ *Emx1*^*IRES-Cre/IRES-Cre*^ (strain number 005628), *Rosa26*^*LSL-tdTomato/LSL-tdTomato*^ (strain number 007909), *Ai65*^*RCFL-tdT/RCFL-tdT*^ (strain number 021875), and *Rosa26*^*LSL-LacZ/LSL-LacZ*^ (strain number 003474) mice were purchased from Jackson Laboratories. *Gsx2-iCre* mice were generated by Prof. Nicoletta Kessaris and colleagues,^36^ and were initially provided by Prof. Kessaris and Prof. David Price, then purchased from Jackson Laboratories (strain number 025806). *Emx1*^*IRES-FlpO/IRES-FlpO*^ mice were generated previously.^8,9^

### Immunocytochemistry

Mice were transcardially perfused with cold PBS, followed by 4% paraformaldehyde (PFA) in PBS, and brains and spinal cords were dissected and postfixed in 4% PFA/PBS at 4ºC overnight. Tissue was sectioned at 50 µm thickness on a vibrating microtome (Leica) or on a cryostat (Leica). Non-specific binding was blocked by incubating tissue and antibodies in 2% donkey serum (Millipore, S30-100ML)/0.3% BSA (Sigma, A3059-100G) in PBS with 0.3% Triton X-100, or 0.3% BSA in PBS with 0.3% Triton X-100. Tissue sections were incubated with primary antibodies at 4°C overnight. Secondary antibodies were chosen from the Alexa series (Invitrogen), and used at a dilution of 1:500 for 3–4 hours at room temperature (RT). For DAPI staining, tissue was mounted in DAPI-Fluoromount-G (SouthernBiotech, 0100-20).

Primary antibodies and dilutions used: rabbit anti-DARPP32 (Chemicon, AB1656, 1:250); mouse anti-GAP43 (Millipore, MAB347, 1:500); rat anti-Bcl11b (Abcam, ab18465, 1:2,000; detects N-terminus); rabbit anti-Bcl11b (Novus, NB100-2600, 1:250; detects C-terminus), rabbit anti-GFP (Invitrogen, A11122, 1:1,000); chicken anti-GFP (Invitrogen, A10262, 1:500); rabbit anti-RFP (Rockland, 600-401-379, 1:500); goat anti-Cdh13 (R&D, AF3264, 1:200); normal goat IgG control (R&D, AB-108-C; 1:1,000); rat anti-L1CAM (Millipore, MAB5272, 1:500); rat anti-Somatostatin (Chemicon, AB5494, 1:100); rabbit anti-NP-Y (Immunostar, 22940, 1:500); mouse anti-Parvalbumin (Sigma-Aldrich, P3088, 1:500); and rabbit anti-b-galactosidase (Rockland, 200-4136, 1:500).

### Nissl staining

Tissue was initially prepared as for immunocytochemistry above, then mounted on gelatin-coated slides and allowed to dry. Tissue was wetted, then stained in 0.25% cresyl violet for 4 minutes. Tissue was rinsed and dehydrated through a series of ethanol and xylene baths (Two minutes each: 50% EtOH, 70% EtOH, 95% EtOH, 100% EtOH; 45 minutes: xylenes), and mounted in DPX.

### Anterograde labeling

Projection neurons were labeled by insertion of crystalline 1,1′.dioctadecyl-3,3,3′,3′-tetramethylindocarbocyanine perchlorate (DiI) into sensorimotor cortex of P0 fixed tissue. Tissue was incubated in PBS/0.05% sodium azide at 37 °C for four weeks to allow DiI transport to occur. DiI-labeled brains were sectioned sagittally at 70 µm on a vibrating microtome. Images shown are a composite of multiple sagittal montages collapsed from 3-D into a single projected 2-D image of the corticospinal tract.

### Retrograde labeling

Developmental SCPN at P0 or P1 were retrogradely labeled unilaterally from rostral brainstem by injecting 690 nl of the retrograde label Green RetroBeads^TM^ IX (Lumafluor) guided by ultrasound backscatter microscopy (VisualSonics, Vevo 3100) via a pulled glass micropipette (Drummond Scientific, 3-000-203-G/X) with a digitally-controlled, volume-displacement nanojector (Nanoject II, Drummond Scientific). For these neonatal injections, pups were deeply anesthetized by hypothermia under ice for 4 minutes. After injections, pups were placed on a heating pad at 37 ºC for recovery.

CSN at P21 were retrogradely labeled bilaterally from the cervical C1 segment by CTB-555 (2 mg/ml in PBS; Thermo Scientific, C22843). For these P21 injections, mice were anesthetized with isoflurane. After laminectomy at the C1 vertebral segment, 322 nl of CTB-555 solution was injected into each side of the midline via a pulled glass micropipette with a Nanoject II. The skin was sutured, and mice were placed on a heating pad at 37ºC for recovery. Mice were subsequently perfused at P25 for analysis of CTB-labeled CSN in cortex.

### DNA constructs

The EGFP-GPI expression construct was derived by subcloning cDNA encoding *Acrosin* signal peptide:hemagglutinin-tag:EGFP:GPI attachment signal peptide into a pCBIG vector backbone, a gift from C. Lois. *Cdh13* cDNA purchased from Sino Biological was subcloned into Cre-activated plasmid backbone (CB-FLEX). For construction of gRNA expression plasmids, the oligonucleotides were inserted into pSPgRNA plasmid, a gift from C. Gersbach (Addgene plasmid # 47108; RRID:Addgene_47108).^86^ Target specific gRNA sequences are as follows: *Cdh13* gRNA#1, 5’-AAAACGAGGGAGCGTTATGA-3’; *Cdh13* gRNA#2, 5’-CAACTTCCCAAATAGATCAG-3’; *Cdh13* gRNA#3, 5’-GGACCAATGGCTTTACAAGA-3’; *Cdh13* gRNA#4, 5’-AATCCGTCTTGTAAAGCCAT-3’; and *LacZ* gRNA, 5’-TGCGAATACGCCCACGCGAT-3’. SP-dCas9-VPR was a gift from G. Church (Addgene plasmid #63798; RRID:Addgene_63798).^65^ dCas9-KRAB-MeCP2 was a gift from A. Chavez & G. Church (Addgene plasmid #110821; RRID:Addgene_110821).^66^

### *In utero* electroporation

Following deep anesthesia of timed pregnant mice, the uterine horns were exposed, and the plasmids, mixed with Fast Green dye (Sigma), were microinjected into one of the lateral ventricles. For electroporation at E14.5, four 35 volt pulses (50 ms on and 950 ms off per pulse) were delivered in appropriate orientation across the embryonic head using 5 mm diameter platinum plate electrodes (CUY650-P5, Protech International Inc) and a CUY21EDIT square wave electroporator (Nepa Gene, Japan). For electroporation at E12.5, four 25 volt pulses (30 ms on and 970 ms off per pulse) were delivered. In the DNA mixture, the concentration was 2 or 4 µg/µl for plasmids expressing cytosolic EGFP or EGFP-GPI, respectively, for labeling CPN for slFSPS. For CRISPR experiments, the concentration was 1 µg/µl for cytosolic EGFP expression plasmid, 1 µg/µl for Cas9 variant expression plasmids, and 2 µg/µl for gRNA expression plasmids. For dual population mosaic labeling (BEAM experiments),^68^ the plasmid concentrations were as follows: 50 ng/µl for the Cre-expression plasmid, 0.5 µg/µl for the CreM plasmid (Cre-inducible Cre recombinase), 2 µg/µl for the Cre-activated myristoylated tdTomato plasmid, 2 µg/µl for the Cre-activated Cdh13 plasmid, and 2 µg/µl for the Cre-silenced myristoylated mClover3 plasmid.

### Soma purification (FACS)

Fluorescently labeled SCPN somata were purified according to established approaches.^16,30,44^ Briefly, retrogradely labeled cortical hemispheres were collected in chilled dissociation solution, then micro-dissected in chilled HBSS using a fluorescence dissection microscope to remove meninges and ventricular zone, thus enrich for labeled SCPN. Cortical tissue was transferred to dissociation solution and enzymatically dissociated for FACS. Fluorescently labeled somata were purifeid from the bulk soma prep by sorting on a customized FACS SORP AriaII instrument equipped with an 85 µm nozzle operating at 45 p.s.i. Somata were sorted directly into the RLT plus buffer (QIAGEN RNeasy Plus Micro Kit) supplemented with 2-Mercaptoethanol for total RNA extraction.

### GC purification (FSPS)

Fluorescently labeled SCPN or CPN axonal GCs were isolated and purified as previously described.^30,31^ Briefly, brainstem and spinal cord were dissected rapidly for SCPN GCs, and cortical contralateral hemisphere was dissected rapidly for CPN GCs. Dissected tissues were homogenized in 0.32 M sucrose supplemented with 4 mM HEPES (Thermo Fisher, 15630106), Halt protease and phosphatase inhibitor cocktail (Thermo Fisher, 78442), and RNasin ribonuclease inhibitors (Promega, N2515) at 900 RPM in a glass-Teflon tissue homogenize tube with 10-12 strokes. Following a low speed centrifugation at 1,660 *g* for 15 min at 4 °C, “post-nuclear homogenate” (PNH) supernatant was collected, and layered onto a 0.83 M and 2.5 M sucrose gradient (Top-to-bottom: PNH, 0.83 M sucrose, 2.5 M sucrose). The discontinuous sucrose gradient was centrifuged at 242,000 *g* for 47 min at 4 °C in a vertical rotor (VTi50 rotor, Beckman Coulter). The bulk GC fraction (GCF) was extracted from the interface between 0.32 M and 0.83 M sucrose.

Labeled axonal GCs were purified from bulk GCF via fluorescent small particle sorting (FSPS), utilizing a customized BD SORP FACS AriaII instrument.^30,31^ Bulk GCF was diluted 4- to 6-fold in chilled PBS prior to loading onto the sorter. GCs were sorted directly into the RLT plus buffer (QIAGEN RNeasy Plus Micro Kit) supplemented with 2-Mercaptoethanol for total RNA extraction.

### GC surface labeling fluorescent small particle sorting (slFSPS)

200 µl CPN GCF was diluted 5-fold in chilled PBS with 3% BSA, then incubated with anti-GFP (Invitrogen, A31852, 1:500), anti-NCAM1 (R&D, FAB7820R, 1:500), or anti-rabbit IgG (1:500) antibodies conjugated to Alexa647 for 60 min on ice. 200 µl SCPN GCF was diluted 5-fold in chilled PBS with 3% BSA, then incubated with normal goat IgG control (R&D, AB-108-C; 1:500), anti-Cdh13 antibody (1:100), or anti-L1CAM antibody (1:500) for 60 min on ice, followed by incubation with a Alexa488-conjugated secondary antibody (1:1,000) for 20 min on ice. Following incubation, GCF was diluted 20-fold with 0.32 M sucrose, and spun at 15,000 *g* at 4ºC for 30 min. After carefully removing supernatant, pelleted GC particles were resuspended in 1 mL 0.32 M sucrose, and diluted 5-fold with chilled PBS prior to loading onto the customized BD SORP FACS AriaII instrument for fluorescent small particle analysis.

### Genome-wide RNA-sequencing and bioinformatic analyses

Following total RNA extraction from purified SCPN GCs and somata, RNA was quality controlled and quantified using an Agilent 2100 Bioanalyzer. Purified RNA samples were converted to cDNA using the SMART-seq v4 Ultra Low Input RNA Kit (Takara Bio), and cDNA libraries were generated with the Nextera XT DNA Library Preparation Kit (Takara Bio) by the Harvard University Bauer Core Facility. High-throughput sequencing was performed by loading the pooled GC and soma cDNA libraries on an Illumina NovaSeq platform. The raw FASTQ files of 100 bp paired-end reads were collected for downstream analysis. Five *Bcl11b* WT and five KO biological replicates were analyzed in both soma and GC analyses.

Adapter sequences were clipped, and unpaired reads were trimmed with Trimmomatic version 0.39. Reads were aligned to the GRCm39 mouse genome with STAR,^87^ and reads mapping to exons were quantified with featureCounts.^88^ Based on read count distributions, genes that had a mean TPM greater than 3 across all replicates, and more than 3 TPM in at least 4 replicates of at least one genotype, were retained for downstream analyses in both soma and GC datasets. To identify “ambient” transcripts among GC transcripts that might arise from supplementing labeled B6 brainstems and spinal cords with unlabeled CD1 cortices during GC purification, variants were called on all samples with GATK HapotypeCaller.^29,89^ 3,492 genes were filtered from differential analysis, since their CD1 penetrance was significantly higher (q < 0.05, one-sided binomial test) than 30% in at least 4 of the 10 GC samples. 7,946 genes passed this quality control test, and were evaluated for differential abundance. Differentially expressed genes between *Bcl11b* WT and KO somata were identified with DESeq2, using an FDR of 0.1.^90^ 10,975 genes had a normalized mean count of >= 23, the independent filtering threshold, and effect sizes were shrunken with the apeglm method.^91^ Since some GC samples clustered by replicate number (Supplementary Figure 5B), potentially due to batch effects, differentially abundant genes between *Bcl11b* WT and KO GCs were identified by DESeq2 pairwise comparison, using “batch” in the experiment design, and an FDR of 0.1. 7,794 genes had a normalized mean count of >= 19, the independent filtering threshold. GO enrichment analysis of biological processes was performed with ClusterProfiler,^92^ and terms were reduced based on semantic similarity with Revigo.^93^ All data are deposited in NCBI’s Gene Expression Omnibus, and will be accessible at GEO Series accession number GSE260678 (https://www.ncbi.nlm.nih.gov/geo/query/acc.cgi?acc=GSE260678) upon publication.

### Imaging

For epifluorescence microscopy, tissue sections and cells were imaged using either a Nikon Eclipse 90i or NiE microscope (Nikon Instruments) with a mounted CCD or sCMOS camera (ANDOR Technology), respectively. Z stacks were collapsed using the “Extended Depth of Focus” function on the NIS-Elements acquisition software (Nikon Instruments). Images were processed using Fiji.^94^ For confocal imaging, samples were imaged on an LSM 780 (Zeiss).

### Cell culture and Immunoblot analysis

HEK 293T cells were cultured at 37ºC with 5% CO^2^ in Dulbecco’s modified Eagle medium (Gibco, 10566024) supplemented with 10% (v/v) fetal bovine serum, 100 U/ml penicillin, and 100 µg/ml streptomycin (Thermo Scientific, 15140122). Cells were transfected using FuGENE HD transfection reagent (Promega, E2311).

One day following transfection, cells were washed with PBS, and lysed with a cell lysis buffer containing 50 mM Tris-HCl (pH 7.5), 150 mM NaCl, 1.0% Triton X-100, 1 mM dithiothreitol, 1 mM EDTA, and Halt protease and phosphatase inhibitor cocktail (Thermo Scientific, 78440). The lysate was centrifuged at 20,000*g* at 4 ºC for 15 min to collect supernatant. Following standard Tris-glycine SDS–PAGE, resolved proteins were electroblotted onto PVDF membranes using semi-dry transfer. Membranes were incubated with primary antibodies diluted in 5% BSA in TBS with 0.1% Tween-20. The following primary antibodies were used for immunoblotting: mouse anti-β-actin (Santa Cruz, sc-47778, 1:1,000); goat anti-Cdh13 (R&D, AF3264, 1:200). HRP-conjugated secondary antibodies were used for ECL imaging. Immunoreactive bands were detected by chemiluminescence using SuperSignal West Pico PLUS (Thermo Scientific, 34580), which was visualized using a CCD camera imager (FluoroChemM, Protein Simple).

### Statistics

Statistical analyses were performed using Graphpad Prism 9.0 software. Sample sizes were predetermined by consistency with established studies. A two-tailed t-test or Mann-Whitney test was used for comparison of two data sets. A two-way ANOVA followed by Bonferroni multiple comparison test was used for experiments to investigate progressive SCPN axon extension. Equal variances between groups and normal distribution of data were presumed, but not formally tested.

## SUPPLEMENTARY FIGURES

**Supplementary Figure 1.**
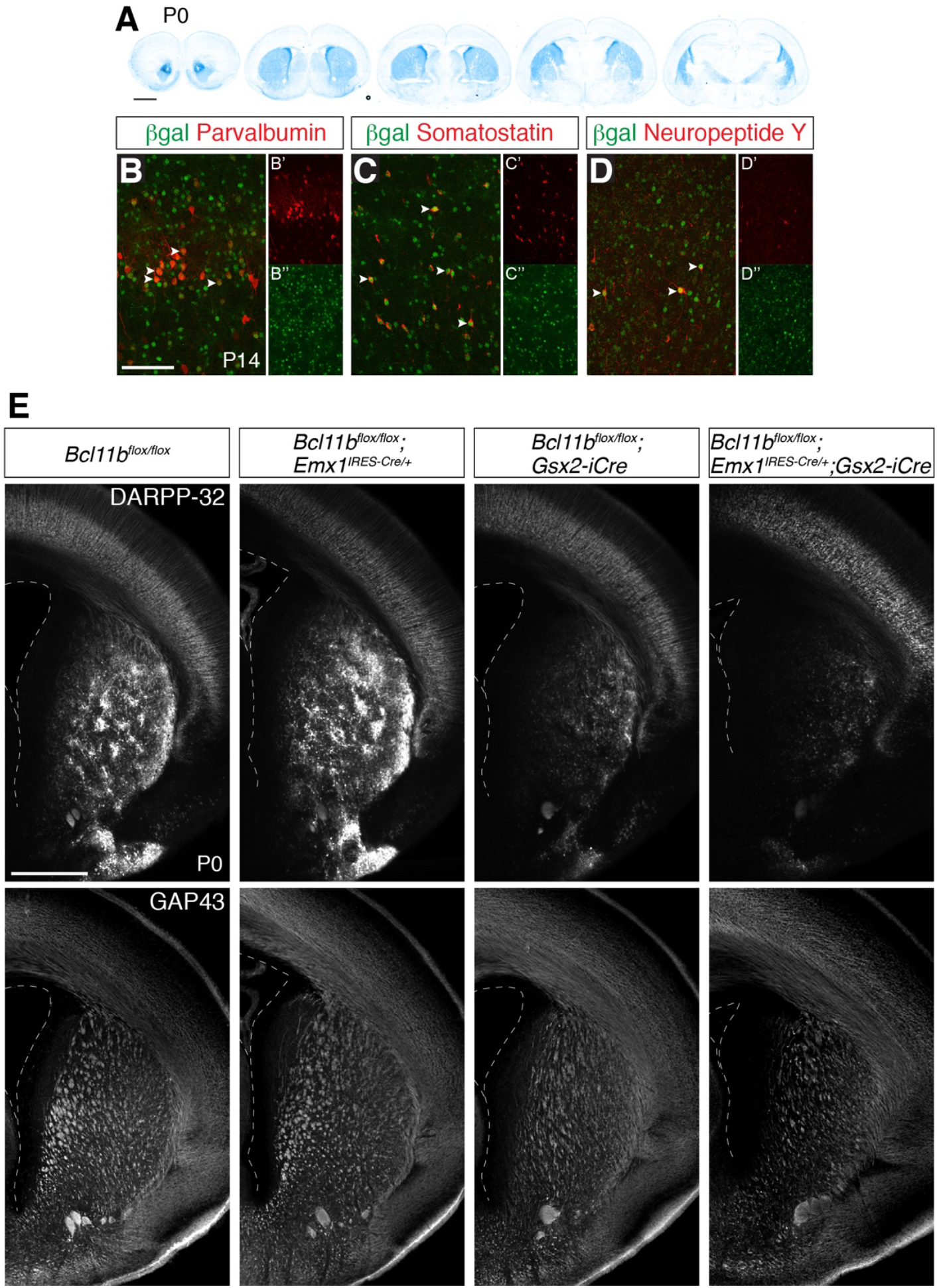
Gsx2-iCre is expressed by neurons derived from the lateral and medial ganglionic eminences. (**A**) In P0 *Gsx2-iCre;Rosa26*^*LSL-LacZ/+*^ forebrains, β-galactosidase is detected in striatum by LacZ labeling. Scale bar, 1 mm. (**B**– **D**) In P14 *Gsx2-iCre;Rosa26*^*LSL-LacZ/+*^ cortex, β-galactosidase co-localizes with parvalbumin, somatostatin, and neuropeptide Y, markers of subpallial-derived interneurons (arrowheads). Scale bar, 100 µm. (**E**) DARPP-32 is expressed normally in *Bcl11b*^*flox/flox*^*;Emx1*^*IRES-Cre/+*^ striatum, but is substantially reduced in *Bcl11b*^*flox/flox*^*;Gsx2-iCre* and *Bcl11b*^*flox/flox*^*;Emx1*^*IRES-Cre/+*^*;Gsx2-iCre* striatum. GAP43 immunolabeling reveals that fasciculation of CFuPN axons in the internal capsule is largely normal in *Bcl11b*^*flox/flox*^*;Emx1*^*IRES-Cre/+*^ striatum, but is substantially disrupted in *Bcl11b*^*flox/flox*^*;Gsx2-iCre* and *Bcl11b*^*flox/flox*^*;Emx1*^*IRES-Cre/+*^*;Gsx2-iCre* striatum. Scale bar, 500 µm.

**Supplementary Figure 2.**
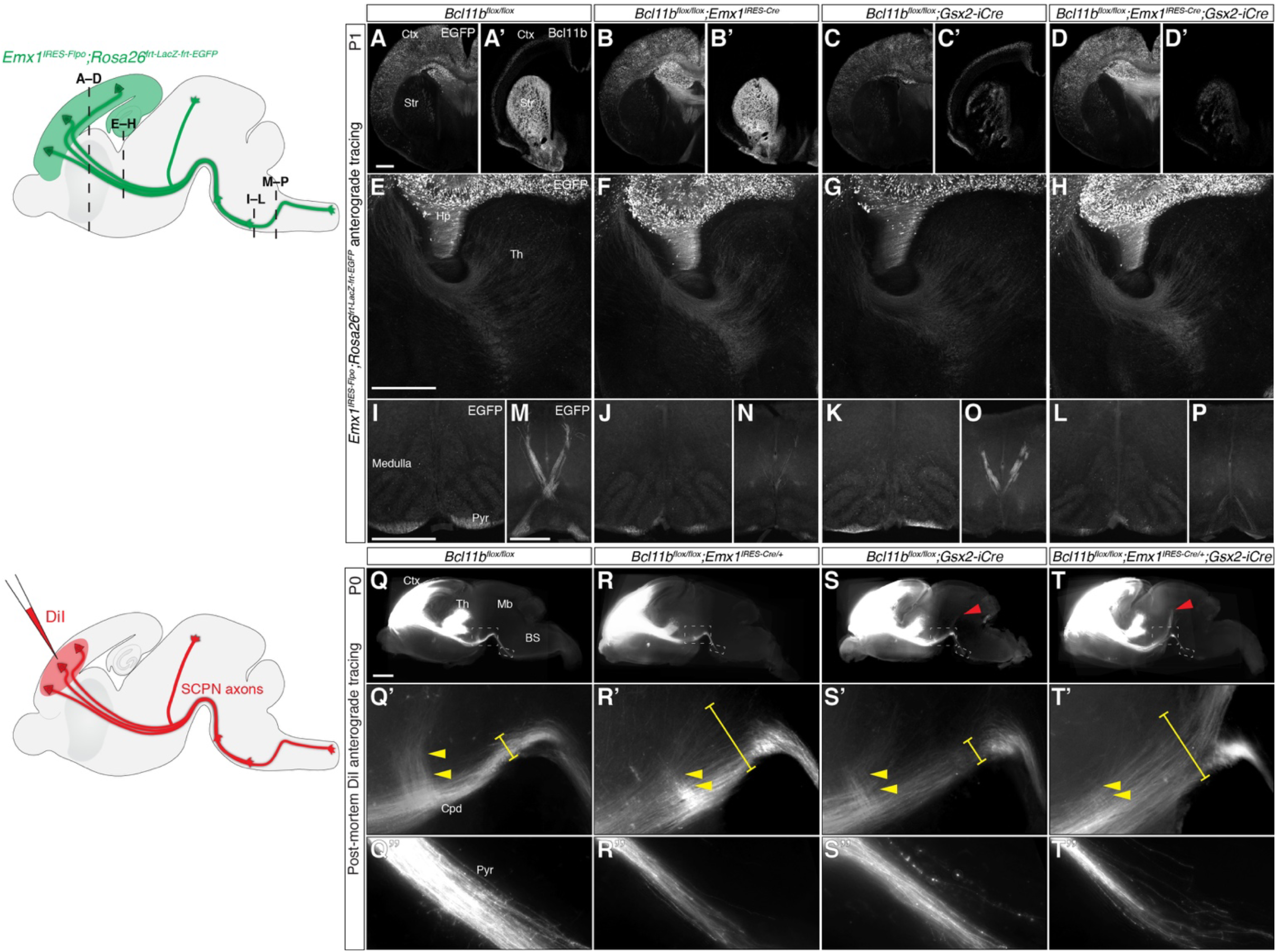
*Bcl11b* deletion from SCPN disrupts outgrowth of SCPN axons, and *Bcl11b* deletion from MSN causes abnormal SCPN axon targeting into dorsocaudal midbrain. (**A**–**P**) Cortical efferent axons are visualized anterogradely by *Emx1*^*IRES-Flpo*^*;Rosa26*^*frt-LacZ-frt-EGFP*^ in coronal sections of P1 brains. Schematic illustrates positions of coronal sections. (A–D) Cortical efferent axons are visualized anterogradely by *Emx1*^*IRES-Flpo*^*;Rosa26*^*frt-LacZ-frt-EGFP*^ in coronal sections of P1 brains. EGFP expression in *Emx1*^*IRES-Flpo/+*^*;Rosa26*^*frt-LacZ-frt-EGFP/+*^ cortex is indistinguishable between wild-type and any of the *Bcl11b* mutant cortices. (A’–D’) Bcl11b expression in cortex and striatum is ablated by *Emx1*^*IRES-Cre*^ and *Gsx2-iCre*, respectively. (E–H) Innervation of thalamus by CThPN axons is largely unchanged in any of the mutant mice compared to wild-type mice. Compared to wild type (I, M), EGFP-labeled SCPN axons reaching the medulla and decussating are substantially reduced in *Bcl11b*^*flox/flox*^*;Emx1*^*IRES-Cre/+*^ mice (J, N), and are slightly reduced in *Bcl11b*^*flox/flox*^*;Gsx2-iCre* mice (K, O). (L, P) Deletion of *Bcl11b* from both cortex and striatum additively reduces SCPN axons reaching the medulla and decussating. Scale bars, 500 µm. (**Q**–**T**) DiI crystals placed in sensorimotor cortex of fixed P0 brains anterogradely label efferent axons, including SCPN axons, as schematized in a sagittal view. (Q–Q’’) DiI labeled axons reaching caudal to thalamus are unequivocally SCPN axons. (Q’) In wild-type (*Bcl11b*^*flox/flox*^) brains, DiI labeled axons fasciculate tightly in the cerebral peduncle (bracket). (Q, Q’’) While some SCPN axons (i.e., corticotectal axons) branch and turn dorsally to enter the lenticular fascicle (yellow arrowheads), others extend further caudally into pons, and leading SCPN axons reach caudal brainstem. (R–R’’) In *Bcl11b*^*flox/flox*^*;Emx1*^*IRES-Cre/+*^ brains, SCPN axons lacking *Bcl11b* fail to fasciculate tightly in the midbrain, corticotectal axons do not turn precisely to enter the lenticular fascicle (R’, bracket, yellow arrowheads), fewer SCPN axons reach rostral brainstem, and only a few leading axons reach caudal brainstem. (S–S’’) In *Bcl11b*^*flox/flox*^*;Gsx2-iCre* brains, SCPN axons branch normally and turn toward tectum, and fasciculate largely normally in the cerebral peduncle (S’, bracket, yellow arrowheads). However, a small subset of labeled SCPN axons abnormally target into dorsocaudal midbrain (S, red arrowheads), and there are slightly fewer SCPN axons reaching caudal brainstem (S’’). (T–T’’) SCPN axons in brains lacking *Bcl11b* in both cortex and striatum are more severely affected than either single mutant. SCPN axons are defasciculated in the cerebral peduncle (T’, bracket), do not branch toward tectum (T’, yellow arrowheads), abnormally target dorsocaudal midbrain (T, red arrowheads), and do not appropriately reach caudal brainstem (T, T’’). Scale bar, 1 mm. BS, brainstem. Cpd, cerebral peduncle. Ctx, cortex. Str, striatum. Hp, hippocampus. Pyr, pyramidal tract. Th, thalamus.

**Supplementary Figure 3.**
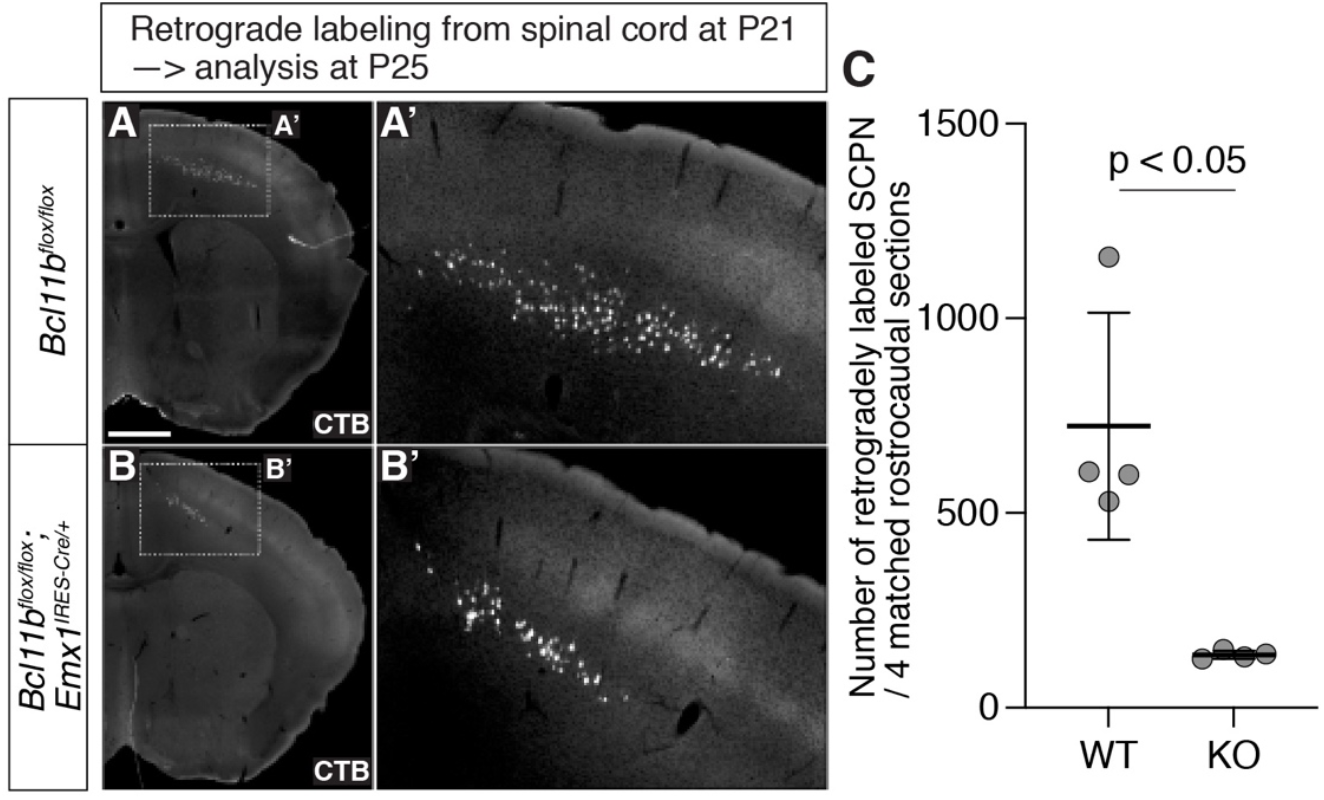
*Bcl11b* deletion in cortex substantially reduces corticospinal projections. (**A, B**) Fewer SCPN are retrogradely labeled in P25 *Bcl11b*^*flox/flox*^*;Emx1*^*IRES-Cre/+*^ cortex by Alexa-conjugated cholera toxin B (CTB) injection into the cervical spinal cord at P21. (**C**) Quantification of retrogradely labeled neurons reveals that loss of *Bcl11b* reduces the number of SCPN axons reaching the spinal cord. Mean ± SEM, n = 4. Mann-Whitney test. *; p < 0.05. Scale bar, 1 mm.

**Supplementary Figure 4.**
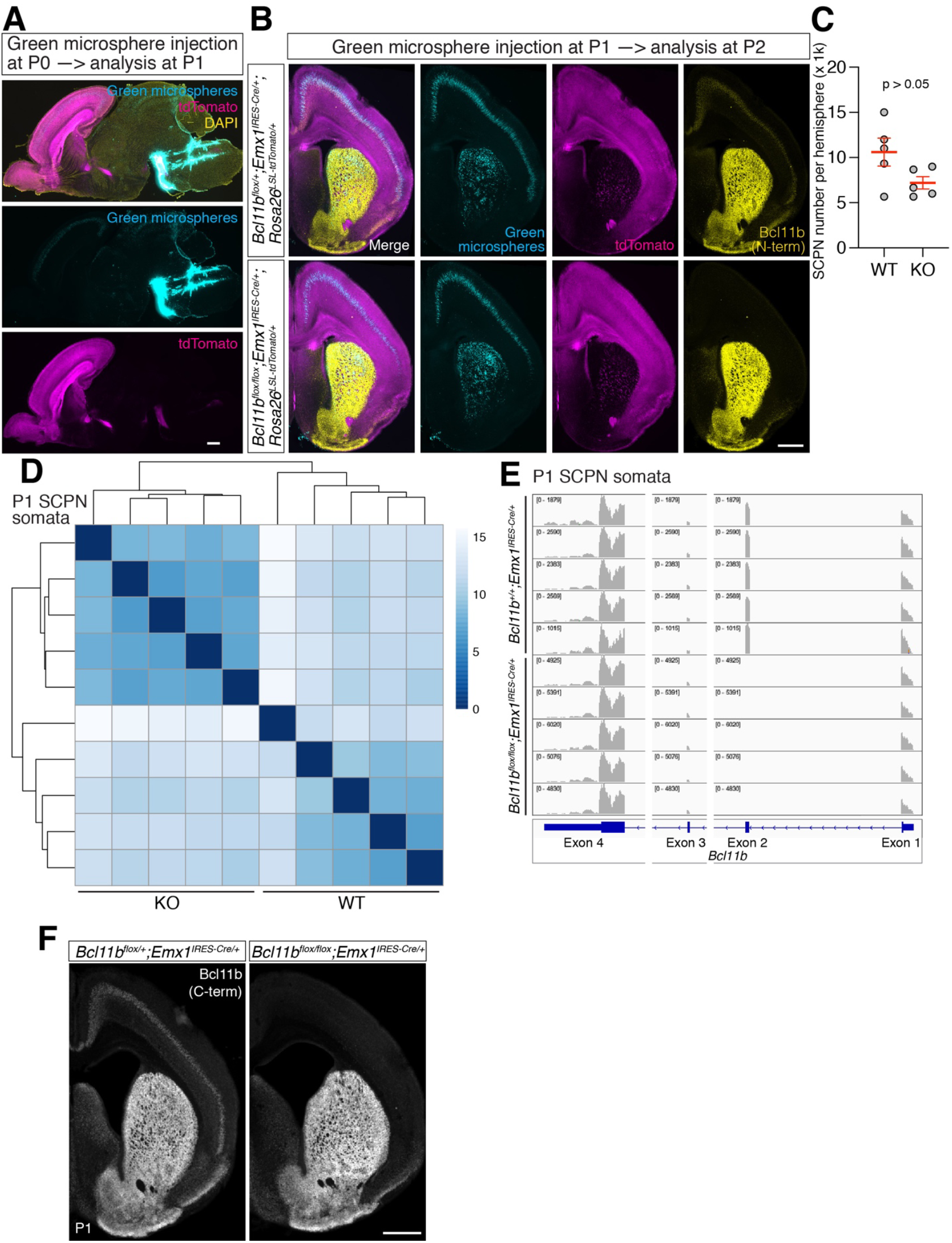
Purification of SCPN somata from wild-type and cortex-specific *Bcl11b* null mice. (**A**) A representative P1 brain image showing green-fluorescent microsphere (cyan) injection into the rostral brainstem of a P0 *Emx1-Cre;Rosa26tdTomato* mouse, and retrogradely labeled SCPN in cortex. The micropipet was inserted rostrally from caudal brainstem. tdTomato, magenta. DAPI, yellow. (**B**) Distribution of retrogradely labeled SCPN is largely normal in *Bcl11b*^*flox/flox*^*;Emx1*^*IRES-Cre/+*^ cortex, with minor reduction of retrogradely labeled SCPN in lateral cortex. Green-fluorescent microspheres (cyan) were injected into P1 *Emx1-Cre;Rosa26tdTomato* mouse, and brains were fixed at P2. tdTomato, magenta. Bcl11b (N-terminus), yellow. (**C**) FACS-isolated SCPN (tdTomato^+^ and green microsphere^+^) per hemisphere is not significantly reduced by *Bcl11b* deletion from cortex. Green fluorescent microspheres were injected at P0, and SCPN were collected at P1. Mean ± SEM, n = 5. Student’s t-test. (**D**) Clustering of soma samples by genotype, using the 500 most variable genes. Lower sample-to-sample distances have darker colors, and higher sample-to-sample distances have lighter colors. WT, SCPN somata from *Bcl11b*^*+/+*^*;Emx1*^*IRES-Cre/+*^*;Rosa26-tdTomato*^*flox/+*^ mice. KO, SCPN somata from *Bcl11b*^*flox/flox*^*;Emx1*^*IRES-Cre/+*^*;Rosa26-tdTomato*^*flox/+*^ mice. (**E**) Exon 2 of *Bcl11b* is deleted in SCPN of *Bcl11b*^*flox/flox*^*;Emx1*^*IRES-Cre/+*^*;Rosa26-tdTomato*^*flox/+*^ mice. (**F**) Efficient deletion of Bcl11b in cortex of P1 *Bcl11b*^*flox/flox*^*;Emx1*^*IRES-Cre/+*^ mice, demonstrated by an antibody detecting the C-terminus of Bcl11b (encoded by exon 4). Scale bars, 500 µm.

**Supplementary Figure 5.**
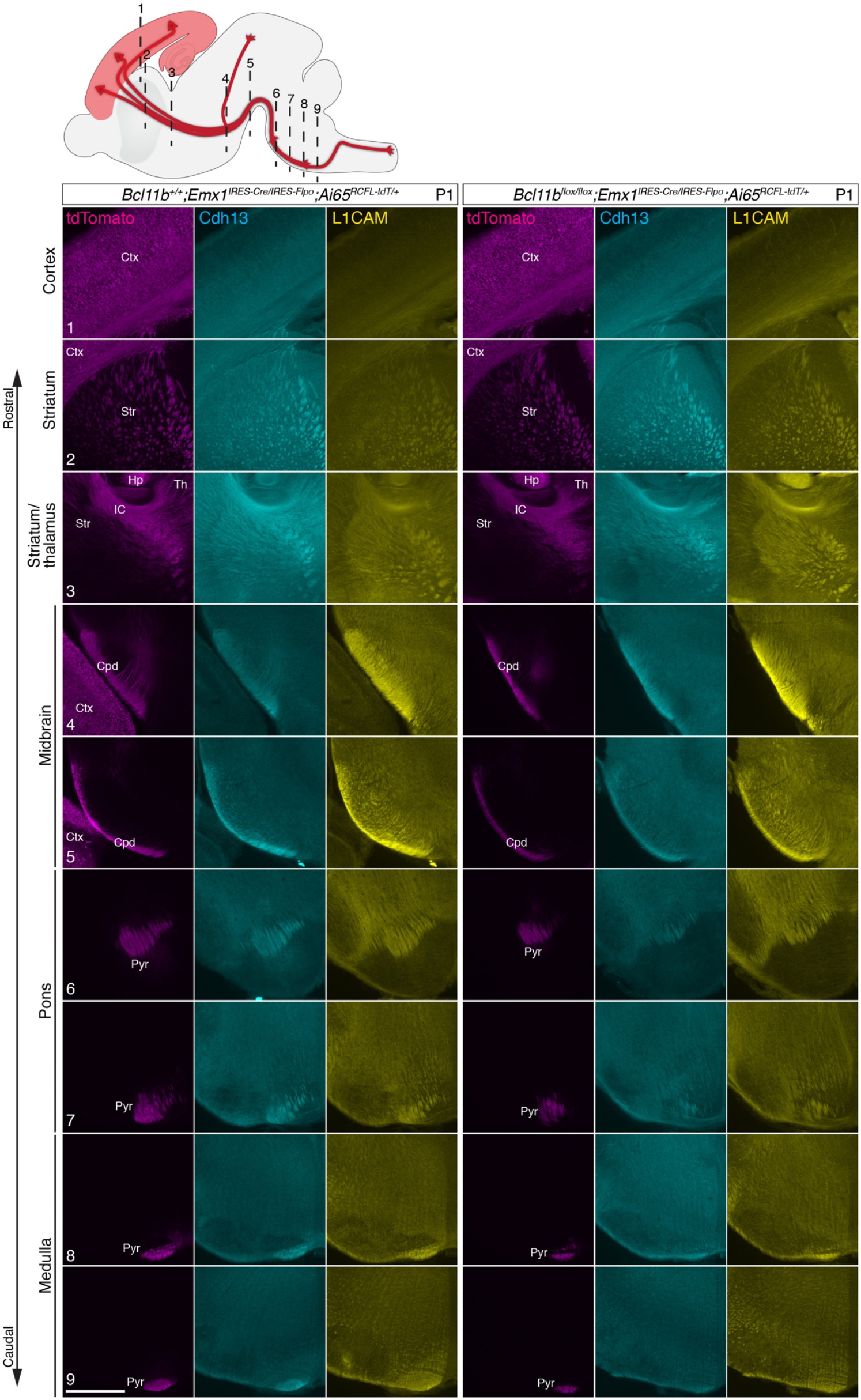
Cdh13 abundance along SCPN axons decreases after *Bcl11b* deletion in cortex. Schematic illustrates positions of coronal sections. Immunocytochemistry of Cdh13 (cyan) and L1CAM (yellow), and tdTomato fluorescence (labeling *Emx-*expressing cell lineage, magenta) of P1 coronal brain sections. After deletion of *Bcl11b* from cortex, fewer tdTomato-positive SCPN axons reach brainstem, and Cdh13 abundance along SCPN axons is substantially reduced in brainstem. Note that reduction in L1CAM abundance along SCPN axons is proportional to the reduction in SCPN axons after *Bcl11b* deletion from cortex. Scale bar, 500 µm. Cpd, cerebral peduncle. Ctx, cortex. Hp, hippocampus. IC, internal capsule. Pyr, pyramidal tract. Str, striatum.Th, thalamus.

**Supplementary Figure 6.**
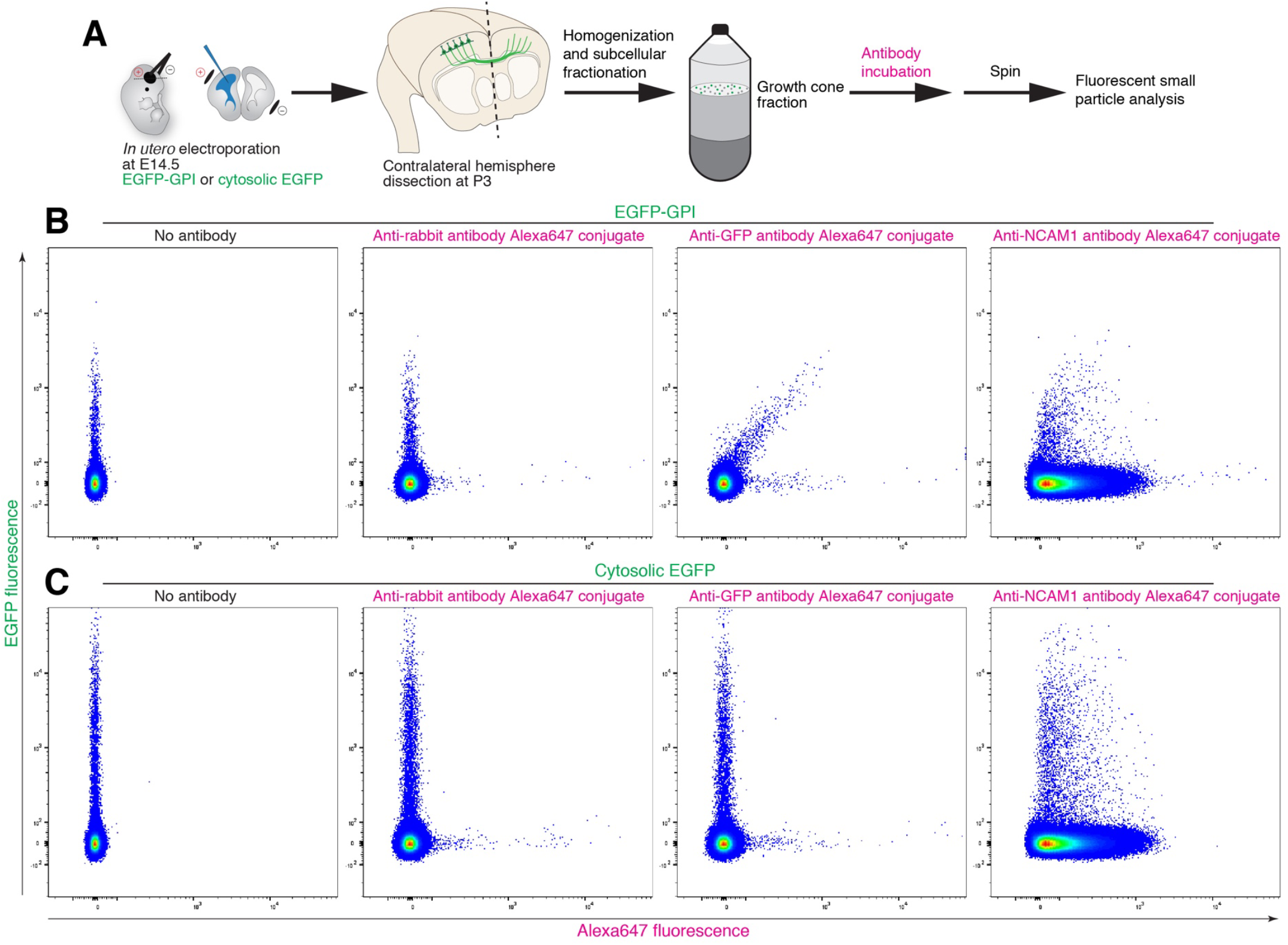
Growth cone surface labeling enables quantitative investigation of antigens on GC surfaces (slFSPS). (**A**) Schematic illustrating slFSPS to investigate antigens on GC surfaces. Here, we labeled CPN unilaterally by *in utero* electroporation at E14.5 with a plasmid encoding EGFP-GPI or cytosolic EGFP, and isolated bulk GCs from the contralateral hemisphere at P3 by subcellular fractionation. After incubation with an antibody conjugated to Alexa647, we diluted and centrifuged GC fractions to remove unbound antibodies, and performed FSPS. (**B**) EGFP-GPI CPN GCs are labeled by anti-GFP and anti-NCAM1 antibodies. (**C**) Cytosolic EGFP CPN GCs are labeled by anti-NCAM1 antibody, but not by anti-GFP antibody.

**Supplementary Figure 7.**
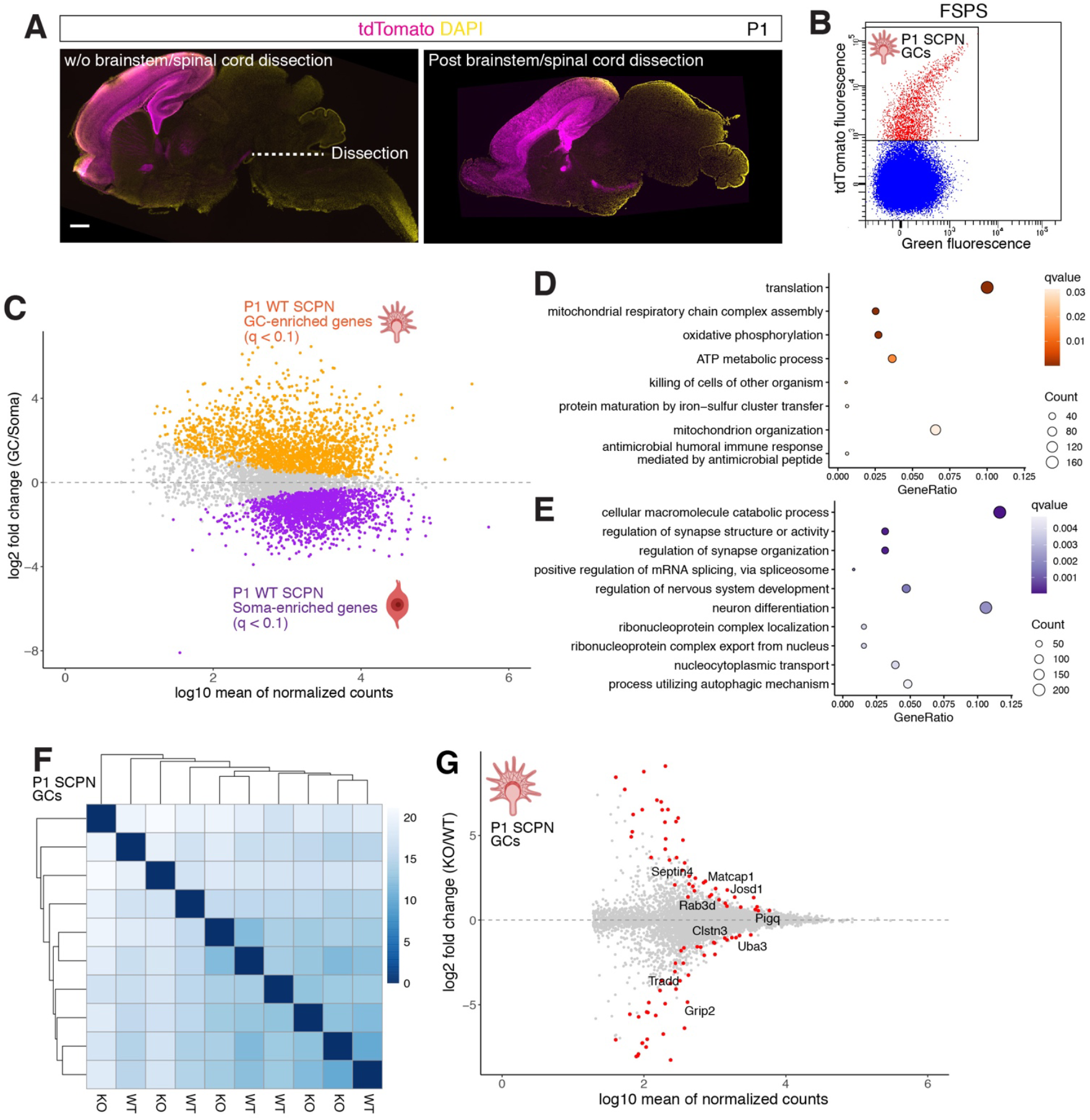
SCPN GC purification and subcellular transcriptomic analyses. (**A**) Representative images showing dissection of brainstem and spinal cord. Upper panel shows a sagittal section of P1 *Emx1-Cre;Rosa26tdTomato* CNS tissue without brainstem/spinal cord dissection. Lower panel shows a sagittal section of P1 *Emx1-Cre;Rosa26tdTomato* CNS tissue after brainstem/spinal cord dissection. tdTomato, magenta. DAPI, yellow. Scale bar, 500 µm. (**B**) A representative FSPS plot is shown. (**C**–**E**) (C) Differences in transcript abundance in *Bcl11b*^*+/+*^*;Emx1*^*IRES-Cre/+*^ (WT) SCPN GCs vs. somata. 2,049 GC-enriched genes (q < 0.1) are shown in orange, and 1,938 soma-enriched (q < 0.1) are shown in purple. Other genes are shown in gray. Total: 6,396 genes. These compartments show highly distinct transcriptomes, highlighting tightly regulated transport of these transcripts along SCPN axons. (D, E) GO enrichment analysis of GC-enriched (D) and soma-enriched (E) genes in *Bcl11b*^*+/+*^*;Emx1*^*IRES-Cre/+*^ SCPN. All (D) and the 10 most significantly enriched (E) terms are shown. Points are colored by q-value, and sized by number of genes per term. GC-enriched genes (q < 0.1) are over-represented for GO terms associated with translation and mitochondrial functions (see also Supplementary Table 9), consistent with previous findings from distinct neuronal populations,^31,62^ suggesting that GCs across distinct neuronal subtypes have partially shared transcriptomic characteristics. Soma-enriched genes (q < 0.1) are over-represented for GO terms associated with neuronal differentiation and functions (see also Supplementary Table 9), reflecting SCPN soma processes at P1. Together, these findings demonstrate successful SCPN GC purification. (**F, G**) (F) Clustering of GC samples, using the 500 most variable genes. Lower sample-to-sample distances have darker colors, and higher sample-to-sample distances have lighter colors. WT, SCPN GC from *Bcl11b*^*+/+*^*;Emx1*^*IRES-Cre/+*^*;Rosa26-tdTomato*^*flox/+*^ mice. KO, SCPN GC from *Bcl11b*^*flox/flox*^*;Emx1*^*IRES-Cre/+*^*;Rosa26-tdTomato*^*flox/+*^ mice. (G) Changes to transcript abundance in SCPN GCs upon deletion of *Bcl11b* by *Emx1-Cre*. 88 differentially abundant genes (DAGs; q < 0.1) are shown in red. Other genes are shown in gray. Total: 7,794 genes. DAGs include genes relevant to cytoskeleton (*Septin4, 49314228F04Rik* (recently named *Matcap1*)), trafficking (*Rab3d, Grip2*), signaling (*Tradd*), protein modification (*Josd1, Uba3*), glycosylphosphatidylinositol (GPI)-anchor biosynthesis (*Pigq*), and adhesion (*Clstn3*).

**Supplementary Figure 8.**
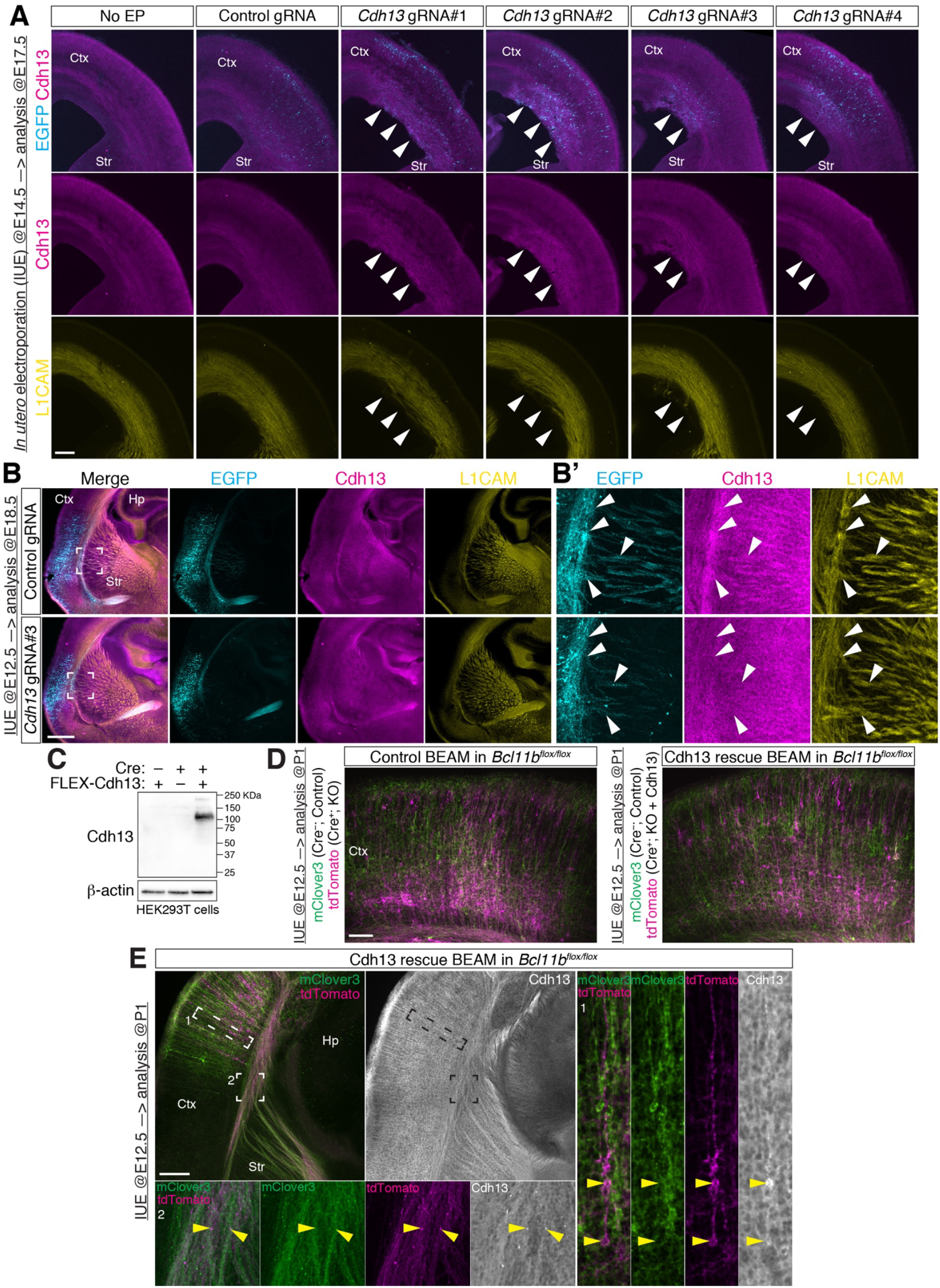
Manipulation of Cdh13 expression *in vivo*. (**A**) Fluorescence images of E17.5 wild-type brains electroporated *in utero* at E14.5 with a plasmid encoding the indicated gRNA, together with those encoding EGFP and dCas9-VPR. A littermate brain without electroporation is shown as a negative control. All 4 *Cdh13* gRNAs induce ectopic Cdh13 expression (arrowheads), while control gRNA does not. None of the gRNAs alter L1CAM expression. EGFP, cyan. Cdh13, magenta. L1CAM, yellow. Scale bar, 200 µm. (**B**) Fluorescence images of E18.5 wild-type brains electroporated *in utero* at E12.5 with a plasmid encoding control gRNA or *Cdh13* gRNA#3, together with those encoding EGFP and dCas9-KRAB-MeCP2. Areas in the inset are enlarged in B’. EGFP-labeled axons in striatum derived from neurons expressing control gRNA colocalize with Cdh13 and L1CAM (B’, arrowheads). EGFP-labeled axons in striatum derived from neurons expressing *Cdh13* gRNA#3 exhibit reduced Cdh13, and unchanged L1CAM immunolabeling (B’, arrowheads). EGFP, cyan. Cdh13, magenta. L1CAM, yellow. Scale bar, 500 µm. (**C**) Immunoblot analysis of Cdh13 overexpression in HEK293T cells. Cells were transfected with plasmids encoding Cre recombinase and/or Cre-activated Cdh13 (FLEX-Cdh13), then cultured for 1 day. (**D**) Fluorescence images of P1 *Bcl11b*^*flox/flox*^ brains electroporated *in utero* at E12.5 in control BEAM or Cdh13 rescue BEAM experiments. Fluorescently labeled cells in the cortex are shown. mClover3, green. tdTomato, magenta. Scale bar, 100 µm. (**E**) Fluorescence images of P1 *Bcl11b*^*flox/flox*^ brains electroporated *in utero* at E12.5 in Cdh13 rescue BEAM experiments. Fluorescently labeled cells and axons in the forebrain are shown. Cdh13 abundance is increased in Cre^+^ tdTomato-labeled somata and axons (insets 1 and 2, yellow arrowheads). mClover3, green. tdTomato, magenta. Cdh13, white. Scale bar, 200 µm. Ctx, cortex. Str, striatum. Hp, hippocampus.

## SUPPLEMENTARY TABLES

**Supplementary Table 1. Comparison between *Bcl11b***^***+/+***^***;Emx1***^***IRES-Cre/+***^ **and *Bcl11b***^***flox/flox***^***;Emx1***^***IRES-Cre/+***^ **SCPN soma transcriptomes**.

List of all SCPN soma genes that were analyzed by DESeq2, and their expression changes after *Bcl11b* deletion. Columns correspond to gene names, means of normalized counts, log2-fold expression changes, standard error estimates (lfcSE), p-values, q-values, and ENSEMBL gene IDs. Positive log2-fold expression changes indicate increased abundance after Bcl11b deletion.

**Supplementary Table 2. Differentially expressed genes in SCPN somata after *Bcl11b* deletion**.

List of 515 SCPN soma DEGs (q < 0.1) after *Bcl11b* deletion. Extracted from Supplementary Table 1.

**Supplementary Table 3. Comparison of transcriptional regulators controlling cortical projection neuron development between *Bcl11b***^***+/+***^***;Emx1***^***IRES-Cre/+***^ **and *Bcl11b***^***flox/flox***^***;Emx1***^***IRES-Cre/+***^ **SCPN somata**.

List of transcriptional regulator genes expressed by P1 SCPN that play key roles in cortical projection neuron development, and their expression changes after *Bcl11b* deletion. Extracted from Supplementary Table 1.

**Supplementary Table 4. Comparison of SCPN marker genes between *Bcl11b***^***+/+***^***;Emx1***^***IRES-Cre/+***^ **and *Bcl11b***^***flox/flox***^***;Emx1***^***IRES-Cre/+***^ **SCPN somata**.

List of SCPN marker genes expressed by P1 SCPN, and their expression changes after *Bcl11b* deletion. Extracted from Supplementary Table 1.

**Supplementary Table 5. GO term enrichment analysis of SCPN soma DEGs after *Bcl11b* deletion from SCPN somata**. List of GO terms that are significantly over-represented (q < 0.05) among 515 SCPN soma DEGs. All SCPN soma genes were used as background. Columns correspond to GO term IDs, descriptions, q-values, gene counts, gene ratio, and gene names.

**Supplementary Table 6. Comparison of axon guidance genes between *Bcl11b***^***+/+***^***;Emx1***^***IRES-Cre/+***^ **and *Bcl11b***^***flox/flox***^***;Emx1***^***IRES-Cre/+***^ **SCPN somata and GCs**.

List of axon guidance genes expressed by P1 SCPN and/or abundant in P1 SCPN GCs, and their expression/abundance changes after *Bcl11b* deletion. Extracted from Supplementary Tables 1 and 8.

**Supplementary Table 7. Comparison between *Bcl11b***^***+/+***^***;Emx1***^***IRES-Cre/+***^ **SCPN soma and GC transcriptomes**.

List of all *Bcl11b*^*+/+*^*;Emx1*^*IRES-Cre/+*^ SCPN soma and GC genes that were analyzed by DESeq2, and their subcellular mRNA localization. Columns correspond to gene names, means of normalized counts, log2-fold transcript abundance changes, standard error estimates (lfcSE), p-values, q-values, and ENSEMBL gene IDs. Positive log2-fold transcript abundance changes indicate higher transcript subcellular localization in GC compartment over soma compartment.

**Supplementary Table 8. SCPN GC transcriptome comparison between *Bcl11b***^***+/+***^***;Emx1***^***IRES-Cre/+***^ **and *Bcl11b***^***flox/flox***^***;Emx1***^***IRES-Cre/+***^.

List of all SCPN GC transcripts that were analyzed by DESeq2, and their abundance changes after *Bcl11b* deletion. Columns correspond to gene names, mean of normalized counts, log2-fold transcript abundance changes, standard error estimates (lfcSE), p-values, q-values, and ENSEMBL gene IDs. Positive log2-fold transcript abundance changes indicate increased abundance after Bcl11b deletion.

**Supplementary Table 9. GO term enrichment analysis of GC-enriched and soma-enriched genes in *Bcl11b***^***+/+***^***;Emx1***^***IRES-Cre/+***^ **SCPN**.

List of GO terms that are significantly over-represented (q < 0.05) among 2,049 GC-enriched and 1,938 soma-enriched genes in *Bcl11b*^*+/+*^*;Emx1*^*IRES-Cre/+*^ SCPN. All shared genes between SCPN GC and soma genes (6,396 genes) were used as background. Columns correspond to GO term IDs, descriptions, q-values, gene counts, gene ratio, enriched compartment, and gene names.

**Supplementary Table 10. Differentially abundant genes in SCPN GCs in *Bcl11b* deletion**.

List of 88 SCPN GC DAGs (q < 0.1) after *Bcl11b* deletion. Extracted from Supplementary Table 8.

